# Spatiotemporal analysis of SARS-CoV-2 infection reveals an expansive wave of monocyte-derived macrophages associated with vascular damage and virus clearance in hamster lungs

**DOI:** 10.1101/2023.03.22.533759

**Authors:** Ola Bagato, Anne Balkema-Buschmann, Daniel Todt, Saskia Weber, André Gömer, Bingqian Qu, Csaba Miskey, Zoltan Ivics, Thomas C. Mettenleiter, Stefan Finke, Richard J. P. Brown, Angele Breithaupt, Dmitry S. Ushakov

## Abstract

Factors of the innate immune response to SARS-CoV-2 in the lungs are pivotal for the ability of the host to deal with the infection. In humans, excessive macrophage infiltration is associated with disease severity. Using 3D spatiotemporal analysis of optically cleared hamster lung slices in combination with virological, immunohistochemical and RNA sequence analyses, we visualized the spread of SARS-CoV-2 through the lungs and the rapid anti-viral response in infected lung epithelial cells, followed by a wave of monocyte-derived macrophage (MDM) infiltration and virus elimination from the tissue. These SARS-CoV-2 induced innate immune processes are closely related to the onset of necrotizing inflammatory and consecutive remodelling responses in the lungs, which manifests as extensive cell death, vascular damage, thrombosis, and cell proliferation. Here we show that MDM are directly linked to virus clearance, and appear in connection with tissue injury and blood vessel damage. Rapid initiation of prothrombotic factor upregulation, tissue repair and alveolar cell proliferation results in tissue remodelling, which is followed by fibrosis development despite a decrease in inflammatory and anti-viral activities. Thus, although the hamsters are able to resolve the infection following the MDM influx and repair lung tissue integrity, longer-term alterations of the lung tissues arise as a result of concurrent tissue damage and regeneration processes.

## Introduction

Infection with Severe Acute Respiratory Syndrome Coronavirus 2 (SARS-CoV-2) can lead to acute respiratory distress syndrome in humans, which is associated with lung injury and pneumonia with high mortality rates. Despite successful vaccination programmes and an increasing prevalence of less virulent SARS-CoV-2 variants, there are still many cases where infection leads to severe respiratory disease. Epithelial cells of upper and lower respiratory tracts express receptors for SARS-CoV-2 cell entry, rendering these tissues the primary infection sites in humans. It is now well established that the characteristics of early immune responses to SARS-CoV-2 are associated with severity of the disease [1]. Essential to this are myeloid lineage cells, which play a crucial role in this first response in both human and animal SARS-CoV-2 infection [2–4]. In healthy lungs, tissue-resident alveolar macrophage and more heterogeneous interstitial macrophage populations are distinguished by their origin, molecular marker repertoire and association with specialized tissue structures [5–7]. The macrophage population composition changes dramatically upon lung infection by respiratory viruses [6, 8, 9]. Loss of tissue-resident macrophages leads to infiltration of bone marrow-derived monocytes in the airway niche [6, 8]. The abundance of myeloid cells in COVID-19 airways has been linked to high levels of myeloid chemoattractants [10], and severe disease development is associated with chemokine receptor gene control in monocytes and macrophages [11]. Moreover, the clearance of dead or dying cells impairs the anti-inflammatory function of macrophages [12].

More detailed investigations, such as temporal omics analysis of SARS-CoV-2 infected hamster lungs have characterized infected lung cell populations and demonstrated extensive changes in innate immune profiles over time [13]. However, despite the wealth of data obtained since the start of the COVID-19 pandemic, studies of pulmonary immune response to SARS-CoV-2 in humans and animal models provided only limited spatial information. With respect to other tissues, such as the upper respiratory tract, information is even more limited. On the other hand, delineating the timing and spatial tissue distribution of immune cells following viral infection is essential for understanding the early immune response, and it may have wider implications since this process is not unique to SARS-CoV-2 infection, but occurs upon inflammation caused by a number of respiratory viruses [14, 15].

Previously, we have shown that following the orotracheal infection of Golden Syrian hamsters, the peak of the clinical signs and virus shedding were already reached between 5 and 7 days post infection (dpi) [16]. This model is therefore ideal to study the early immune response upon SARS-CoV-2 infection within the first 7 days. Studies using the hamster SARS-CoV-2 infection model demonstrated that it largely phenocopies the moderate form of COVID-19 in humans, inducing focal diffuse alveolar destruction, hyaline membrane formation, and mononuclear cell infiltration, making it an important tool for detailed studies of tissue infection and the subsequent immune response [13, 17–21]. The hamster model is particularly reminiscent of human COVID-19 in showing strong and age dependent lung infection [19], and virus replication in hamster lungs as well as its pathological effects are reduced when infected with the less pathogenic omicron (B.1.1.529) variant [22].

Moreover, a transient increase in monocyte-derived macrophage (MDM) level in lungs was shown by single-cell RNA-seq analysis [13], reflecting the CD68^+^ cell infiltration that was reported in human COVID-19 cases [23–25]. Monocyte infiltration into tissues following viral infections is well described. These cells differentiate and acquire functional macrophage characteristics already in the blood stream [26], and play a crucial role in shaping the response to infection [27]. Nevertheless, the association of MDM with disease outcome is mainly based on their presence in severely affected tissues, but it remains unclear if MDM directly contribute to immunopathology, because there is still no data tracking the spatial distribution of MDM over time.

To understand the mechanisms of virus entry into tissues, progress to severe pathology or virus clearance at the tissue level, it is necessary to characterize virus localization and spread in specialized tissue compartments and changes in tissue cellular composition, including local distribution of immune cells in relation to patterns of infection at different stages after its initiation. Advanced imaging techniques such as Light-sheet fluorescence microscopy (LSFM) of optically clear tissue samples offer a way to address these questions by visualizing large tissue volumes at cellular resolution. To date LSFM has been applied in three studies visualizing SARS-CoV-2 infection in different animal models [28–30]. Zaeck et al [28] used the ferret model for valuable insights into infection initiation in the epithelium of the upper respiratory tract. The hamster model was used by Tomris et al [29] to demonstrate virus localization in lungs with respect to ACE2 and TMPRSS2 expression, which is a prerequisite for virus entry. Finally, Nudell et al [30] developed a novel technique to visualize SARS-CoV-2 in the entire body of K18-hACE2 transgenic mice. Here we applied a combination of 3D imaging, virological, histopathological and transcriptomic analyses to reveal spatiotemporal characteristics of SARS-CoV-2 infection and MDM infiltration associated with virus clearance, as well as tissue damage and regeneration in hamster lungs.

## Results

### Time-course of SARS-CoV-2 infection

In previous studies, we have established the orotracheal infection of hamsters as the most efficient route [16]. Here we applied this approach to obtain more detailed spatiotemporal information about the early immune response and the change within lung tissue following orotracheal hamster inoculation with 1x10^5^ TCID_50_ units of ancestral SARS-CoV-2.

The infected animals developed increasing clinical signs (weight loss, lethargy, respiratory distress) from 2 dpi, which continued until 7 dpi (Suppl. Table 1, Suppl. Fig. 1a). Assessment of viral RNA nasal shedding showed high levels from 1 to 3 dpi (Suppl. Fig. 1b). Accordingly, we also determined high levels of viral RNA in lungs from 1 dpi, which further increased at 2 dpi (p < 0.01), followed by a gradual decrease on the following days (Fig.1a). In contrast, much lower or no viral RNA was detected in other examined tissues (Suppl. Fig. 1c). Furthermore, the highest levels of replication competent virus were detected in the lung tissue at 1 and 2 dpi, which rapidly decreased until 5 dpi. At 6 and 7 dpi, no replicating virus was detectable anymore, although the level of viral RNA only minorly decreased until 7 dpi, and was still detectable at 14 dpi.

Immunohistochemical analysis identified the bronchial epithelium and alveolar epithelial cells to be positive for SARS-CoV-2 nucleoprotein (NP) from 1 dpi until 7 dpi. At 14 dpi, viral antigen was no longer detected. The highest semi-quantitative NP score was obtained for 1 dpi (Fig. 1b). The NP distribution pattern changed over time and shifted from the large bronchi to the bronchioles and alveolar epithelium (Fig. 1c). Mainly ciliated bronchial epithelial cells and type 2 pneumocytes were affected, and, to a lesser extent, non-ciliated bronchial cells and type 1 pneumocytes. (Fig. 1c).

**Figure 1.**
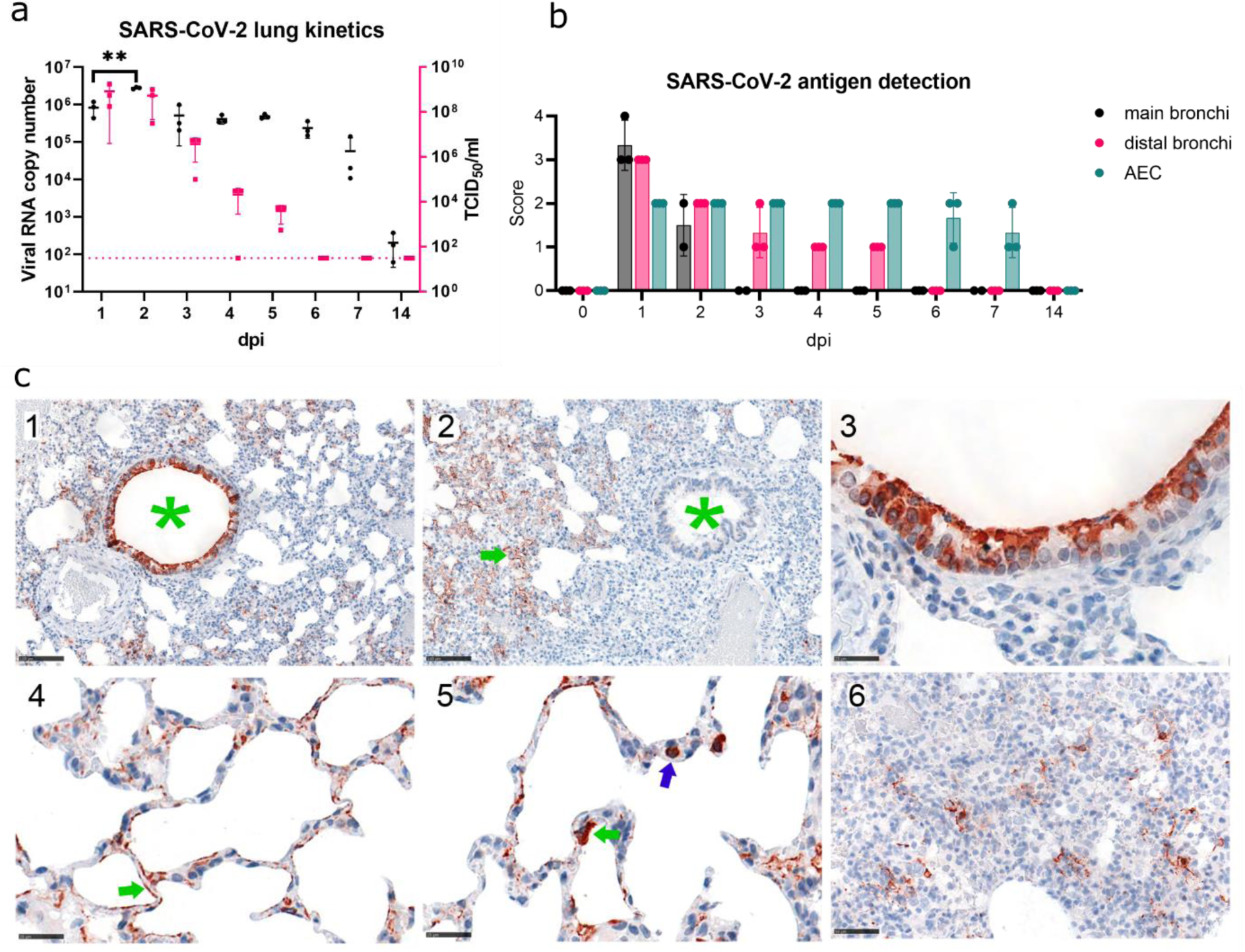
Virological and histopathological characterization of SARS-CoV-2 lung infection time-course. a) SARS-CoV-2 kinetics in hamster lungs. Viral RNA copy number (Log_10_) (black, left axis) and replication competent virus (Log_10_ TCID_50_/ml; magenta, right axis) detected in the lung samples. Each point represents the data from approximately 1.5 mm^3^ sample from right cranial lobe of one animal. The dotted line indicates the detection limit for replication competent virus. b) Immunohistochemical analysis of SARS-CoV-2 NP antigen distribution in lung tissue. Viral antigen score in main bronchi, distal bronchi and alveolar epithelial cells (AEC), control or 1 – 14 days post infection (dpi). Dots, individual animals; bar, median. Brown - NP. Scores correspond to 0 = no antigen, 1 = rare, <5% antigen labeling per slide; 2 = multifocal, 6%–40%; 3 = coalescing, 41%–80%; 4 = diffuse, >80%. c) 1 - Viral antigen distribution at 1 dpi showing strong labelling in bronchi (green asterisk) and AEC. Bar 100 µm. 2 - Viral antigen distribution at 5 dpi, showing labelling restricted to AEC (green arrow), no labelling in bronchi (green asterisk). Bar 100 µm. 3 - Viral antigen in cells morphologically consistent with ciliated bronchial epithelium at 1dpi. Bar 25µm. 4 - Viral antigen in cells morphological consistent with type 2 pneumocytes (green arrow) at 1dpi. Bar 25 µm. 5 - Target cell, viral antigen in cells morphologically consistent with type 1 pneumocytes (green arrow), also intravascular cells (blue arrow) are labelled, interpreted to be monocytes/macrophages taking up viral antigen at 1dpi. Bar 25 µm. 6 - Viral antigen clearing, scattered between proliferating pneumocytes and infiltrating immune cells the antigen is found at 7 dpi. Bar 50µm.

### SARS-CoV-2 lung infection induces a wave of MDM influx

We then applied LSFM to investigate the distribution of infected cells in hamster lungs during the first seven days after inoculation. We also analyzed the infiltration of MDM characterized by anti-CD68 staining using three animals at each time point. Immunostaining and tissue optical clearing of thick lung sections were based on our Ethyl cinnamate (ECi) approach which we previously applied for 3D imaging of SARS-CoV-2 infected ferret tissues [28]. As shown in Suppl Fig 2a, hamster lung tissue transparency was achieved, permitting fluorescence microscopy analysis of SARS-CoV-2 infection, as well as relevant cellular markers.

**Figure 2.**
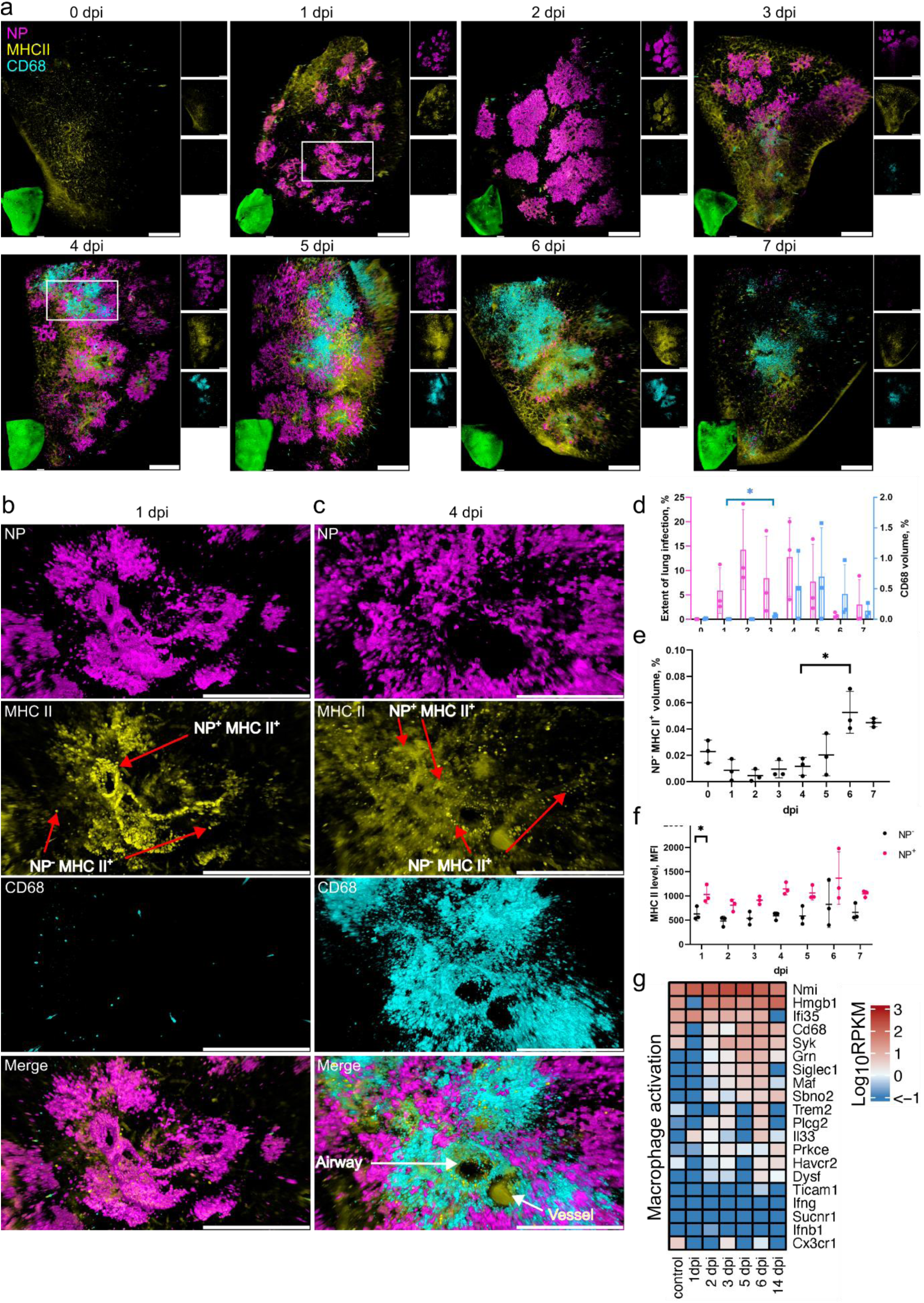
SARS-CoV-2 lung infection leads to an influx of MDM and MHC II upregulation in infected cells. a) LSFM imaging of the infection time-course and corresponding host responses. Lung tissues from three animals for each time point were optically cleared and stained for NP (magenta), MHC II (yellow) and CD68 (cyan). 3D transparency rendering with top view of samples oriented in xy plane at each time point. Large panels show overlays of NP, MHC II and CD68. Bottom left inlay shows tissue autofluorescence (green). Small panels show fluorescence in individual channels: NP (top), MHC II (middle) and CD68 (bottom). Representative images from a single animal are presented. Bar 1000 μm. White rectangles at 1 and 4 dpi correspond to detailed views in b and c, respectively. b) Detailed view of an infected lung region at 1 dpi. c) Detailed view of an infected lung region with a cluster of CD68^+^ cells at 4 dpi. The neighbouring airway and blood vessel were distinguished by autofluorescence. Scale bar 500 μm. d) LSFM quantification showing changes in NP (magenta, left axis) and CD68 (cyan, right axis) levels relative to total tissue volume during post-infection time-course. e) LSFM quantification of NP^-^ MHCII^+^ levels relative to total tissue volume. f) LSFM quantification of MHCII mean fluorescence intensity (MFI) level in tissue regions positive and negative for NP. g) Heat map depicting scaled mRNA expression of genes associated with macrophage activation.

LSFM imaging revealed large nodes of NP staining in bronchial airways already early after infection starting from 1 dpi (Fig. 2a). Despite the relatively limited resolution of the LSFM imaging approach, it showed the presence of CD68 positive immune cells even in uninfected lungs. High-resolution confocal fluorescence microscopy showed that these CD68^+^ cells are also MHC II positive, and are localized in the alveolar compartments, as expected for alveolar macrophages (Suppl. Fig. 2b). Virus spread from the main bronchial epithelial airways to distal regions during the first days post infection was also observed, in agreement with our immunohistochemical data (Fig. 1b) and an earlier hamster study [19].

Our machine-learning based 3D image quantification of LSFM records (Suppl. Movie 1) showed that maximum SARS-CoV-2 levels were reached at 2 dpi, detected in ∼14% of the lung tissue volume (Fig. 2a, d). Analysis of the tissue infection patterns also showed substantial changes at 2 dpi with infection nodes increasing in number and size (Suppl. Fig. 3). The total CD68 volume remained relatively low at 1 and 2 dpi (Fig. 2a, b, d). However, a small but significant (p < 0.05) increase was detected at 3 dpi, suggesting the start of MDM infiltration (Fig. 2a, d). Further substantial changes were observed from day 4, when the initial influx of MDM turned into an MDM wave flooding large regions of the lung (Fig. 2a, c). Notably, the distribution of SARS-CoV-2 infected cells at this time point was uneven, with highly infected but also uninfected regions. At the same time, sites of MDM infiltration did not always coincide with infected regions (Fig. 2c), indicating either a rapid virus clearance in areas of MDM infiltration, or that sites of infiltration are not directly linked to location of infected cells.

The MDM infiltration wave further increased at 5 and 6 dpi, (Fig. 2a, d). In parallel, starting from 4 dpi, the infection nodes also became more fragmented (Suppl. Fig. 3), and by 7 dpi the levels of SARS-CoV-2 NP dropped to less than 5% tissue volume (Fig. 2d). High-resolution confocal analysis confirmed the presence of high-density MDM accumulations, and the proximity of MDM to infected cells, suggesting their direct involvement in virus clearance (Suppl. Fig. 4b).

This observation was further confirmed by immunohistochemistry and quantitative 2D image analysis showing waves of macrophage influx peaking on day 5 (Suppl. Fig. 5). In detail, non-infected control animals showed evenly distributed Allograft Inflammatory Factor 1 (AIF1)-positive macrophages with only single cells found perivascularly (Suppl. Fig. 5b:1a-c). One day after infection, immune cell rolling and perivascular aggregation of monocytes/macrophages occurs (Suppl. Fig. 5b:2a-c). A prominent perivascular cuff comprising many macrophages was present at day 5, with some monocytes/macrophages found “rolling” at the endothelium (Suppl. Fig. 5b:3a-c, b4b-c). At the same time, in areas with high-grade pneumonia, additional massive macrophage aggregates were found in the alveoli (Suppl. Fig. 5:b4a). Finally, on day 14, only single AIF1-positive cells were found perivascularly (Suppl. Fig. 5b:5a-c), even in areas that showed a prominent thickening of the interstitium (Suppl. Fig. 5b:5a-b), which is indicative of previous pneumonia.

To further investigate the MDM infiltration in relation to other early immune response characteristics in the lung, we sequenced RNA of the total lung tissues taken at different time points after infection (Fig. 2g). Analysis of genes related to macrophage activation showed no change or decrease of transcript counts at 1 dpi, but their levels increased in the following days. Most noticeably the levels of *Cd68, Maf and Siglec1* peaked at 5 and 6 dpi in agreement with LSFM imaging data. Among these genes, the increase of *Siglec1* in particular indicates an influx of peripheral monocytes, further supporting the notion that CD68 levels increase, as detected by LSFM imaging at these time points corresponds to MDM infiltration.

### SARS-CoV-2 infection leads to rapid MHC II upregulation in bronchial epithelium

We also followed the MHC II status, since it had previously been reported that MHC II levels are upregulated in lung epithelial cells following respiratory viral infections [31–33]. LSFM analysis of MHC II in lung tissues (Fig. 2a) showed that there was no significant change in NP^-^ MHC II^+^ cell levels at 1-5 dpi (Fig. 2e). However, their levels increased at 6-7 dpi (p < 0.05). In contrast, MHC II level increase (p < 0.05) was detected in infected NP^+^ cells already at 1 dpi (Fig. 2b, f). High-resolution confocal microscopy imaging confirmed that higher MHC II levels were present in the infected cells (Suppl. Fig. 4a). LSFM analysis at later time points showed that all infected cells also exhibited elevated levels of MHC II from 2 dpi onwards (Fig. 2c, f).

The infected cells at 1 dpi primarily belong to the bronchial epithelium, which contains heterogeneous cell populations. We further analysed Club cells, representing a major non-ciliated bronchial epithelial cell population at 1 dpi by co-staining with uteroglobin (UG). Although we observed a large population of Club cells in the hamster airway epithelium defined by high UG level, the majority of Club cells were not infected (Fig. 3a-c), in agreement with our IHC data (Fig. 1c). Moreover, they showed a patchy distribution, and the infected regions of bronchial epithelium were largely devoid of any UG producing cells (Fig. 3b). Image analysis showed that the infected cells in bronchial epithelium had a higher MHC II signal compared to uninfected cells (p < 0.05, Fig. 3c). Of note, the MHC II level in Club cells was also elevated, albeit to a lesser extent, compared to other uninfected cells (p < 0.05). These findings were confirmed by high-resolution confocal imaging, which clearly showed the absence NP staining in Club cells while the neighbouring bronchial epithelial cells were infected, with the MHC II levels increased (Fig. 3d).

**Figure 3.**
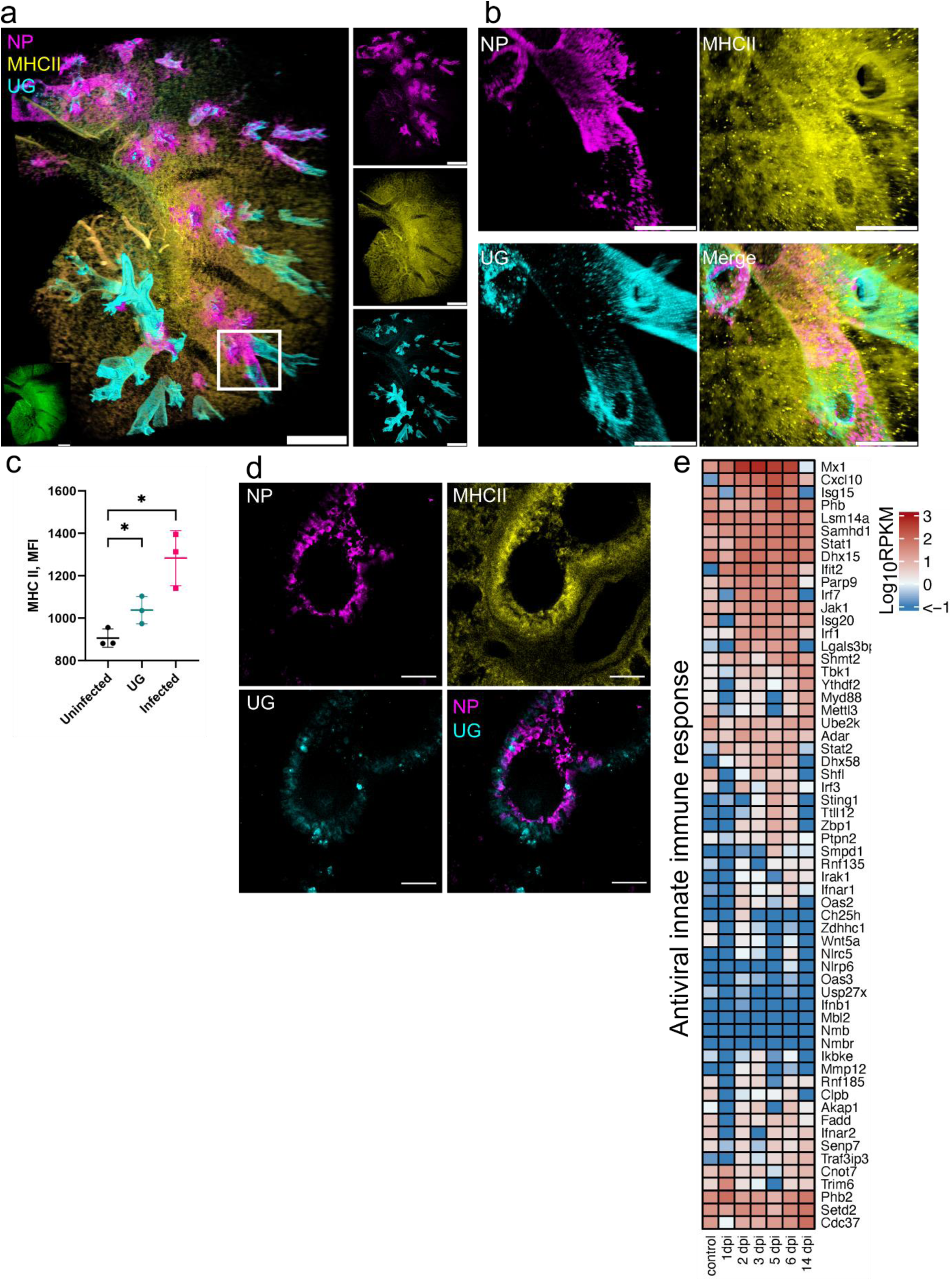
SARS-CoV-2 infection leads to a rapid MHC II upregulation in bronchial epithelium. a) LSFM showing SARS-CoV-2 infection (NP, magenta), MHC II (yellow) and UG staining (cyan) at 1 dpi. 3D transparency rendering with top view of samples oriented in xy plane. White rectangle corresponds to a detailed view of an infected airway region in b. Bottom left inlay shows tissue autofluorescence (green). Small panels show fluorescence in individual channels: NP (top), MHC II (middle) and UG (bottom). Representative images from a single animal are presented. Scale bar 1000 μm. b) Detailed view of an infected airway at 1 dpi. Scale bar 500 μm. c) Quantification of MHC II level (MFI) in UG^-^ uninfected tissue, UG^+^ tissue and UG^-^ infected tissue. d) Confocal imaging of hamster lung at 1 dpi. Maximum intensity projections of NP (magenta), MHC II (yellow), UG (cyan) and a merge of NP and UG. Scale bar 50 μm. e) Heat map depicts scaled mRNA expression of genes associated to type I interferon responses and antiviral immune responses.

To examine the antiviral response, we analysed transcription levels of genes involved in the interferon system and antiviral activities (Fig. 3e). Some of these genes showed little upregulation, or were downregulated at 1 dpi, such as *Ifnb1* and *Irf3*, consistent with previous reports [34]. However, a number of signature genes involved in interferon type I and III, and early viral response, such as *Mx1, Ifit2, Cxcl10, Samhd1* and *Parp9,* were upregulated at 1 dpi, with further increase on the following days (Fig. 3e). Moreover, further upregulation was observed at 2 dpi for another group of genes, including *Irf1, Irf3, Irf7, Isg15* and *Isg20*. Importantly, many of these genes had returned to low levels at 14 dpi (*Mx1, Cxcl10, Ifit2, Irf1, Irf3, Irf7, Isg15*). Nevertheless, many genes remained upregulated or downregulated at 14 dpi compared to uninfected controls.

Increase of STAT1 was recently reported in patients with mild and severe COVID-19 [35]. Remarkably, *Stat1*, *Stat2 and Jak1* were also increased in hamster lungs from 1 and 2 dpi, peaking at 5 to 6 dpi (Fig. 3e). Because STAT1 is required for MHC II induction [36], this suggests that MHC II upregulation is triggered by the canonical Jak-Stat signalling pathway. However, the rapid MHC II increase is not sufficient to completely block virus replication.

### Post-infection endothelial damage is linked to MDM infiltration

Lung vascular hyperpermeability is a major factor contributing to severe COVID-19 [37], which is suggested to result from disruption of the respiratory vascular barrier [38]. The semiquantitative, histopathological evaluation identified high lesion scores at 4-6 dpi (Fig. 4a, b). In detail, vascular lesions and inflammation were detected with particular high scores simultaneously (Suppl. Fig. 6a, b). The identification of necrosis of endothelial cells and cells of the vessel wall, intramural and perivascular inflammation, edema as well as perivascular ferric iron deposition (interpreted as hemosiderin) were indicative for intravital vascular damage and leakage (Fig. 4a, Suppl. Fig. 6c). The earliest evidence for perivascular minimal to mild hemosiderin deposition was found at 4 dpi in 3 out of 3 animals. At 14 dpi, all hamsters showed moderate ferric iron deposits in the lung.

**Figure 4.**
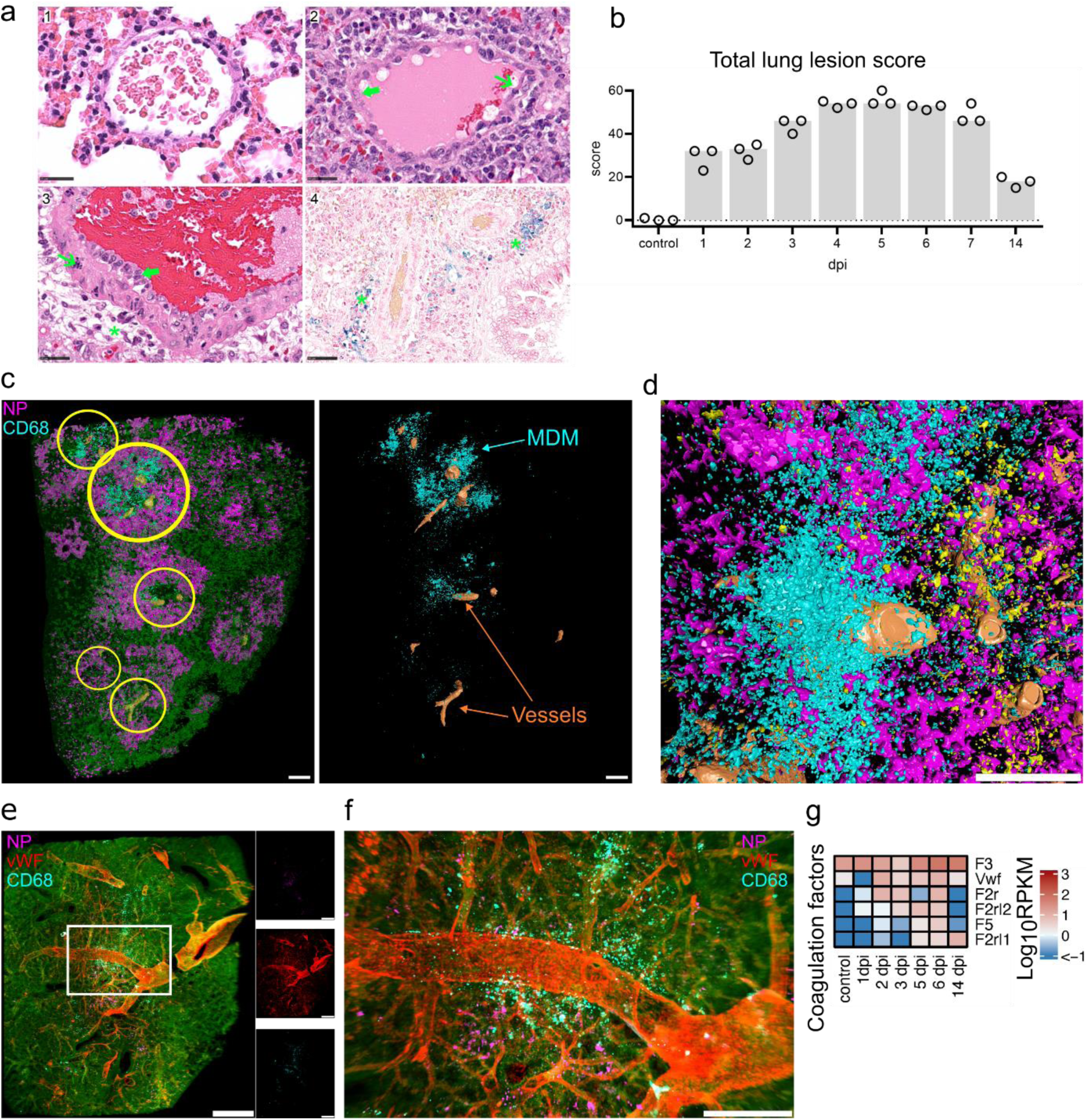
Tissue damage and MDM infiltration. Spatiotemporal analysis of tissue damage and necrosis. a) Exemplary vascular changes in lung tissues in control animals at 6 dpi. (**1**) Normal vein of control animals. (**2**) Vein of infected hamster, 6 dpi showing endothelial hypertrophy (green thick arrow) and necrosis (green slender arrow). (**3**) Artery of infected hamster, 6 dpi with endothelial hypertrophy (green thick arrow), necrosis within the vessel wall (green slender arrow), and perivascular inflammatory infiltrates with edema (green asterisk). (**4**) Blue coloured perivascular (green asterisk) hemosiderin deposition in infected hamsters 6 dpi. Edema and hemosiderin are consistent with vascular leakage. Hematoxylin eosin staining, bar 25 µm (**1-3**) and Prussian blue staining, bar 50 µm (**4**). b) Lung lesion scores. Particularly high scores were found on days 4-6. Total scores were composed of vascular lesion, inflammation, necrosis and regeneration scores (Suppl. Fig. 6). Detailed scoring criteria are given in Suppl. Table 2. c) Machine learning image analysis reveals clusters of infiltrating MDM near major blood vessels. Left: colour coded view of a processed LSFM lung tissue record at 4 dpi showing detection of lung tissue (green), major blood vessels (orange), MDM (cyan) and NP (magenta). Yellow circles indicate MDM clusters juxtaposed to blood vessels. Right: View of MDM and blood vessels only. Bar 500 μm. d) Detailed view of MDM in vicinity of lung vasculature at 5 dpi. Blood vessels (orange), NP (magenta), MDM (cyan) and NP^-^ MHC II^+^ (yellow) objects are shown. Bar 500 μm. e) LSFM imaging of NP and MDM clusters in relation to lung vWF^+^ vasculature at 5 dpi. 3D transparency rendering with top view of samples oriented in xy plane. Large panel shows overlay of NP (magenta), vWF (red) and CD68 (cyan), and tissue autofluorescence (green). Small panels show fluorescence in individual channels: NP (top), vWF (middle) and CD68 (bottom). Representative images from a single animal are presented. Bar 1000 μm. White rectangle corresponds to a detailed view shown on the right. Bar 250 μm. g) Heat map depicts scaled mRNA expression values of coagulation factors associated with extrinsic vessel damage.

**Figure 5.**
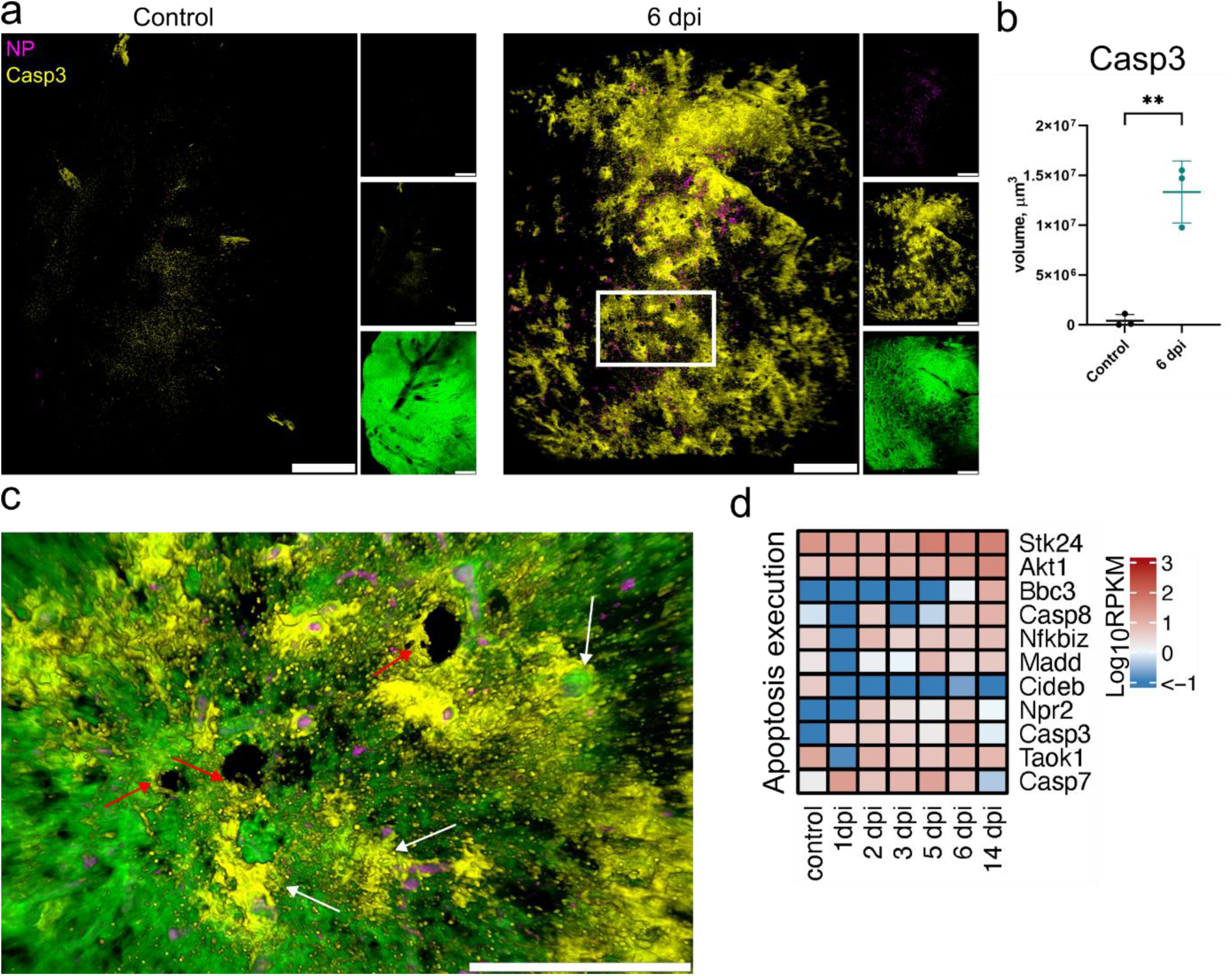
LSFM and transcription analysis of apoptosis following SARS-CoV-2 infection. a) LSFM imaging of cleaved Caspase-3 in hamster lungs in control and at 6 dpi. 3D transparency rendering with top view of samples oriented in xy plane. Large panels show overlays of NP (magenta) and Casp3 (yellow). Small panels show fluorescence in individual channels: NP (top), Casp3 (middle) and tissue autofluorescence (bottom). Representative images from a single animal are presented. Bar 1000 μm. White rectangle corresponds to a detailed view in (c). Scale bar 1000 μm. b) LSFM quantification of Casp3^+^ tissue volume. c) Detailed view of cleaved Casp3 and NP with tissue autofluorescence overlay at 6 dpi. White arrows indicate Casp3 staining near blood vessels, red arrows indicate Casp3 staining near airways. Scale bar 500 μm. d) Heat map depicts scaled mRNA expression of genes associated apoptosis execution

**Figure 6.**
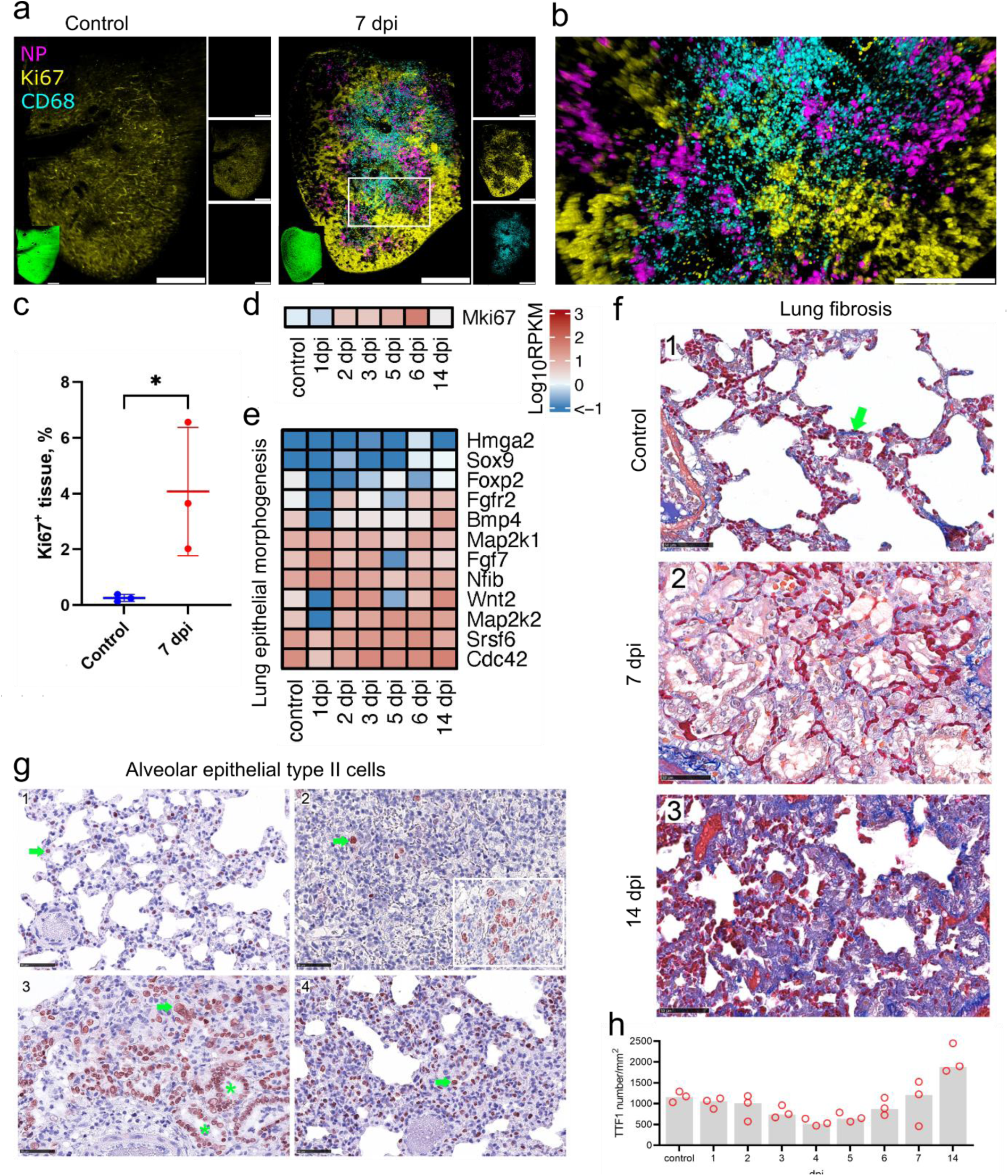
Cell proliferation and fibrosis following SARS-CoV-2 infection. a) Proliferative response visualized by LSFM using Ki67 staining (yellow) in control and at 7 dpi. NP (magenta) and CD68 (cyan) staining were also used to show infection and MDM infiltration. 3D transparency rendering with top view of samples oriented in xy plane. Large panels show overlays of NP (magenta), Ki67 (yellow) and CD68 (cyan). Small panels show fluorescence in individual channels: NP (top), Ki67 (middle) and CD68 (bottom). Representative images from a single animal are presented. White rectangle corresponds to a detailed view in (b). Bar 1000 μm. Detailed view of proliferation at 7 dpi. Bar 500 μm. c) Quantification of Ki67 levels in LSFM records. d-e) Heat maps depict scaled mRNA expression of the Mki67 gene (d) and genes involved in epithelial proliferation in lung morphogenesis (e). f) Azan stain (blue) for collagenous and reticular connective tissue (green arrow). In comparison with non-infected hamsters (**1**), the lungs on day 7 (**2**) show a clearly expanded interstitium, lacking an increase in dark blue collagen fibers. In contrast, 14 days after infection (**3**), the interstitium is expanded, mainly due to an increase in collagen fibers. Azan stain, bar 50 µm. g) Temporal type II pneumocyte (alveolar cell type 2, AT2) detection. Immunohistochemistry (IHC) on TTF1 labelled lung slides. Representative pictures showing IHC for AT2. In control animals (**1**), Day 4 (**2**), day 7 (**3**) and day 14 (**4**) post infection. Pictures showing a red nuclear TTF1 labelling in AT2 cells (green arrow). Note the even distribution of well-differentiated AT2s with small, round nuclei in control animals (**1**). Lung parenchyma affected by virus induced tissue damage shows few TTF1 positive cells (**2**). Oligofocal, there are aggregates of pleomorphic AT2 (**2**, inlay: hypertrophic cells, large nuclei, multinucleated cells) indicating early regeneration. Ongoing regeneration with prominent bronchiolisation of alveoli (green asterisk) by palisading AT2 (**3**). High numbers of AT2 still present at day 14 (**4**), the majority of labelled cells present with small round nuclei indicating later stages of differentiation. Immunohistochemistry, TTF1 antigen, ABC Method, AEC (red) chromogen, hematoxylin counterstain (blue), bar 100 µm (a) and 50 µm (b-, c). g) Quantitative 2D image analysis of TTF1 staining. Following a virus infection induced tissue damage and loss of AT2 on day 3 and 4, AT2 hyperplasia associated increase in relative cell count starts on day 5 and finally exceeds the baseline number on day 14. Dots, relative TTF1-positive number results for individual animals; bar, group median.

Using LSFM image analysis, we were able to identify major blood vessels in lungs based on their autofluorescence. We noted that the infiltrating MDM were often localized in clusters surrounding blood vessels (Fig. 4c, d, Suppl. Movie 1). However, the autofluorescence signal is not sufficient to reliably identify smaller blood vessels and capillaries. To investigate MDM association with vascular structures, we used von Willebrand factor (vWF) as a specific endothelial marker at 5 dpi when both the MDM level and the vascular histopathology scores were high (Suppl. Fig. 6a). Using this approach, we detected characteristic clusters of MDM clearly located near and around the blood vessels (Fig. 4e, f), confirming autofluorescence-based observations. This localization suggests that the MDM may infiltrate in leaky regions following blood vessel injury.

Further transcriptome analysis revealed upregulation of several coagulation factors, with the most increase in tissue thromboplastin (*F3*), *vWF* Coagulation Factor II Thrombin Receptor (*F2r*) and Factor V (*F5*) levels (Fig. 4g). These factors are associated with extrinsic coagulation pathways [39], suggesting the damage of endothelial or subendothelial cells. Moreover, increased transcript numbers of many genes involved in endothelial cell apoptotic process was also detected from 2 dpi and peaked at 6 dpi (Suppl. Fig. 7a).

This data is corroborated by LSFM visualization of cleaved Caspase-3 (Casp3, Fig 5a). This imaging revealed extensive apoptotic regions in lungs at 6 dpi, a time point when the majority of the virus was already cleared (Fig. 2a, d). Quantification of Casp3 positive tissue volume confirmed this observation (p < 0.01, Fig 5b). Noticeably, hotspots of cleaved Caspase-3 were observed near blood vessels and the airways (Fig 5c). Nevertheless, the inflammatory and necrotic processes observed by histopathology suggest that the other cell death mechanisms were also involved. Indeed, post-infection time-course RNA-seq analysis showed an upregulation of *Casp3*, but also a number of other genes involved in apoptosis and necroptosis such as *Stk24, Akt1, Madd, Casp7* and *Casp8* peaking at 5 and 6 dpi (Fig. 4f).

### Lung tissue repair and remodelling following virus clearance

Because the MDM infiltration has previously been linked to proliferation and fibrosis, we analysed the lung proliferation and regeneration status. LSFM analysis of marker of proliferation Ki67 staining at 7 dpi showed regions of proliferative activity (Fig. 6a). A particular spatial distribution pattern of the proliferative upregulation was revealed, with Ki67 being mainly restricted to areas where the viral infection was not detectable (Fig. 6a, b). Moreover, although a large quantity of MDM was observed, the proliferative regions were largely devoid of these cells (Fig. 6a and b). We confirmed the increase in Ki67 level compared to control animals by image analysis (Fig 6c). Further, the *Mki67* transcript level analysis showed the upregulation starting already at 2 dpi and peaking at 6 dpi (Fig. 6d).

The extensive parenchymal and endothelial cell death may lead to compensatory proliferative activity. Indeed, upregulation of *Wnt2*, *Cdc42, Srsf6, Map2k2* and other genes involved in lung epithelial proliferation was detected from 1 dpi (Fig. 6e). Added to this, many genes involved in blood vessel remodelling were upregulated, although this started only at 2 dpi for the majority of these genes (Suppl. Fig. 7b). Most upregulated genes maintained high expression levels even at 14 dpi, as well as the levels of some inflammatory (Fig. 3e), and endothelial apoptotic (Suppl. Fig. 7a) RNA transcripts. Similarly, our histopathological analysis showed that inflammation and regeneration was still detectable at 14 dpi compared to pre-infection levels, although these were lower than at earlier post-infection time points (Suppl. Fig. 6b, d). However, no necrosis was observed at 14 dpi, indicating restoration of tissue integrity (Suppl. Fig. 6c).

We further investigated if hamster SARS-CoV-2 infection induces lung fibrosis. Using azan staining for collagen detection we did not observe any changes at 1-7 dpi compared to control animals (Fig. 6f). However, at 14 dpi an increase in collagen staining and thickening of the interstitium was observed in all hamsters (Fig. 6f). The late detection of fibrosis suggests that collagen accumulation takes several days after induction by MDM or it is not directly caused by the MDM activity.

Regeneration of the conducting airways presents with hypertrophy and hyperplasia of the bronchial epithelium (Suppl. Fig. 8), starting on day 1 post infection (Suppl. Fig. 1d). To determine the regeneration of the alveolar parenchyma, in particular of type 2 pneumocytes (AT2) we used immunohistochemistry and quantitative 2D image analysis.

SARS-CoV-2 infection led to an initial loss of AT2 on days 3 and 4 followed by regenerative AT2 hyperplasia with an increase in relative cell number, starting on day 5, finally exceeding the baseline on day 14 (Fig. 6g, h). In comparison to non-infected control animals (Fig. 6g1), hamsters on 4 dpi showed few remaining AT2 in areas with acute alveolar damage (Fig. 6g2), and pleomorphic and enlarged AT2 were found only in few areas (Fig. 6g2 inlay). Prominent regeneration-associated AT2 hyperplasia presented with bronchiolisation of alveoli and large AT2 nuclei (Fig. 6g3, 7 dpi). Finally, numerous AT2 were still present at day 14 (Fig. 6g4).

Taken together, our data show that in the Golden Syrian hamster model, the proliferative and tissue regeneration activities start as early as 1 dpi in parallel with extensive tissue damage. Importantly, these activities occur simultaneously with the virus spreading and clearance, but continue for a much longer time period after the virus is removed, and the MDM levels are reduced.

## Discussion

The Golden Syrian hamster is generally accepted as a suitable model for the moderate COVID-19 pathogenesis in humans. However, the time course of infection in hamsters is within 7 days, which is distinctly shorter than in humans, where a medium duration of 21 days has been reported [40]. Since the pathology found in hamsters resembles that found in COVID-19 patients, we considered this an ideal model to analyse the spatiotemporal disease progression and antiviral defence in this model.

Combining 3D tissue imaging by LSFM with global transcriptomics and immunohistochemistry revealed signatures of an MDM influx, which coincided with the decrease of SARS-CoV-2 levels. Human autopsy studies showed that severe COVID-19 is associated with profibrotic macrophage response and subsequent proliferative lung fibrosis [23–25]. Similarly, macrophage infiltration was observed in lungs of SARS-CoV-2 infected K18-ACE2 transgenic mice [41] and Golden Syrian hamsters [42, 43]. Here, we observed that MDM appear in large clusters near blood vessels, rather than directly at the sites of infection. The subsequent migration of MDM through the tissue towards the infected cells may be driven by the chemoattractants produced by these cells, as was reported for COVID-19 [10].

Our analysis of SARS-CoV-2 NP localization showed the formation of multiple infection nodes in the epithelial layer of bronchial airways, which rapidly increased in size with virus spreading deeper into the parenchyma. Similar findings were reported in an earlier study showing NP clusters which increased in size during the first days after infection in hamsters [19]. The infection nodes become fragmented by 5-6 dpi with levels of NP decreasing around the bronchi. No new infection nodes were observed, suggesting the absence of virus release into the airways or inhibition of further spread by inflammatory responses. Moreover, we observed selective cell type infectivity, with no infection of the Club cells at 1 dpi. Although we cannot exclude that the lack of detectable infection might be due to a rapid death of infected Club cells, this observation can be explained by the non-ciliated nature of these cells, making them less accessible after virus entry into the airways. This is supported by a recent report that the motile cilia and microvilli are required for virus penetration through the airway mucosal barrier [44].

Substantial lung tissue damage and cell death following SARS-CoV-2 infection have been previously shown in hamsters [13, 23] similar to the lung injury in SARS-CoV-2 infected humans [45]. One of the key consequences is vascular hyperpermeability following disruption of the respiratory vascular barrier [38]. The increased vascular hyperpermeability contributes to severe COVID-19 [37]. vWF and other coagulation factors play a central role in platelet adhesion to damaged vascular subendothelium and cloth formation [46], and their role in thrombosis in COVID-19 was recently shown [47]. Upregulation of coagulation factors associated with endothelial injury suggests that such areas may serve as leaky entry points for MDM. This confirms that the hamster model is also suitable to further analyse the pathomechanisms related to blood clotting. It remains to be verified if the MDM also change their transcriptional profile to a pro-thrombotic signature, similar to that reported in COVID-19 in humans [48]. Taken together, our observations suggest that the vascular hyperpermeability serves as trigger for MDM entry into the tissue and the areas of vascular damage may define the sites of massive MDM infiltration.

Lung tissue contains a variety of MHC II expressing cells, including classical antigen presenting cells and pneumocytes, and MHC II expression in bronchial epithelium is well known [35]. Lung epithelial cells serve as the first line of anti-viral defence by recognizing viral Pathogen-Associated Molecular Patterns (PAMP), which leads to the activation of the immune transcriptional profile and changes in the pulmonary innate immunity [49–51], including recruitment of MDM [9]. One manifestation of alveolar epithelial cell response is the upregulation of MHC II upon respiratory virus infection [52]. Expression of MHC II in human and murine alveolar AT II is well established [31, 53]. It was recently shown that these cells also express the conventional antigen presentation machinery, and thus contribute to improved disease outcome following respiratory viral infections [33]. In the bronchial epithelium, MHC II expression is low in homeostasis, but its level is upregulated in a variety of human lung disorders and upon viral infection [32]. The change in MHC II levels has been suggested to allow immune response modulation, and conversely regulation of the epithelium by the immune system through the release of cytokines by adjacent immune cells [32]. Thus, our observation of MHC II upregulation in infected cells is not surprising and transcriptomics data suggest that this process is triggered by the canonical Jak/Stat signalling pathway. SARS-CoV-2 induces antiviral transcriptional response in both human [54] and hamster [13] lungs. We also observed a broad antiviral response, including induction of interferon type I and III response pathways. The interferon driven response by human epithelial cells is well known, and its robustness is thought to contribute to the severity of COVID-19 [54, 55]. In turn, MHC II associated immune response observed in SARS-CoV-2 infected hamster lungs may lead to changes in barrier integrity, innate immune functions and eventually local cell composition and renewal.

The changes in tissue cellular composition following virus clearance are driven by a number of factors, including immune cell migration, cell death and proliferation. The viral infection inflammatory processes lead to substantial tissue necrosis in hamsters, as shown by our histopathology analysis. Moreover, the transcription levels of genes involved in cell death execution and endothelial apoptosis, as well as increased levels of cleaved Caspase-3 suggest that different cell death mechanisms are taking place, including apoptosis and necroptosis, which have also been shown in earlier SARS-COV-2 studies [56]. On the other hand, the tissue repair processes lead to regeneration of the alveolar epithelial cells and the blood barrier, as clearly seen from IHC, RNAseq and 3D imaging, but also to excessive proliferation of other cells, such as fibroblasts, which leads to collagen deposition and fibrosis, contributing to disease severity [13, 23].

Overall, our data show a rapid anti-viral activity in hamster lungs comprised of a fast antiviral response followed by MDM-driven virus clearance. In parallel, simultaneous tissue damage, thrombotic and proliferative responses occur which are manifested in high levels of apoptosis, vascular damage and an early onset of lung repair. The hamsters are able to resolve most of these processes, including reduction

in myeloid cell levels within two weeks after infection. Nevertheless, some immunopathological characteristics remain, including inflammation and fibrosis. Thus, we propose that the presence of MDM observed late in severe COVID-19 cases may be a consequence of decreased ability to maintain efficient balance between tissue damage and repair, which is critical for complete lung function restoration.

## METHODS

### Animal experiments

### SARS-CoV-2 kinetics

In a previous experiment, we determined the orotracheal inoculation to be the most efficient route to infect hamsters [16]. We therefore chose this route to analyse infection kinetic and host responses within 14 days post infection. Male 5-7 weeks old Golden Syrian hamsters (*Mesocricetus auratus*, raised by Janvier Labs, France) were kept in groups of three to four individuals. Animals were offered water and rodent pellets ad-libitum, fresh hay was offered daily. They were checked for clinical scores and body weight on a daily basis. The hamsters were inoculated orotracheally with 10^5^ TCID_50_ of the ancestral SARS-CoV-2 (isolate 2019_nCoV Muc-IMB-1). To detect viral shedding, nasal wash samples were collected daily under isoflurane anaesthesia by flushing 200 µl PBS along the animal’s nose. Four animals were euthanized by deep isoflurane anaesthesia, cardiac exsanguination and cervical dislocation at days 1, 2 and 3 dpi, while three animals each were sacrificed at days 4 -7. To investigate the disease progression after this time point, another eight animals were sacrificed at 14 dpi. Three mock-infected animals were kept as controls and were sacrificed on day 7 of the experiment. During necropsy, the respiratory tract (nose, trachea, pulmonary lymph nodes, lung), the digestive tract, as well as heart, liver, skeletal muscle and brain were sampled, and aliquots of each tissue were freshly frozen for viral analysis, and the rest stored in 4 % neutral-buffered formalin for histological analysis.

Total RNA was extracted from nasal wash and tissue samples, and SARS-CoV-2 RNA was detected using “Envelope (E)-gene Sarbeco 6-carboxyfluorescein quantitative RT-PCR” as described previously [57, 58].

### Virus titration - tissue culture infectious dose 50 (TCID_50_)

Virus titers were determined as described previously (Blaurock et al., 2022). Briefly, samples were serially diluted in MEM containing 2% FCS and 100 Units Penicillin / 0.1 mg Streptomycin (P/S) (Millipore Sigma, Germany). Vero E6 cells were incubated with 100 µl of ten-fold sample dilutions added in quadruplicates for 1 h at 37°C before 100 µl MEM containing 2% FCS and P/S were added per well and plates were incubated for 5 days at 37°C and 5% CO_2_. Supernatant was removed and cells were fixed with 4% formalin for 30 minutes. Next, the plates were stained with 1% crystal violet for 15 minutes and titers were determined following the Spearman Kaerber method [59].

### Antibodies

The following antibodies used in tissue optical clearing experiments were diluted in 0.5% BSA, 10% DMSO and 0.5% Triton-X-100 in PBS: rabbit anti-SARS-CoV N (Rockland Immunochemicals, #ABIN129544, 1:500), mouse anti-SARS-CoV NP (Sino Biological, #40143-MM05, 1:400), rat anti-I-A/I-E (Biolegend, #107601, 1:400), mouse anti-CD68 (Invitrogen, #MA5-13324, 1:400), rat anti-CD68 (BioLegend, #137001, 1:400), rat anti-Ki-67 (Biolegend, #652401, 1:200), rabbit anti-vWF (Dako, #A0082, 1:1000), rabbit anti-Cleaved Caspase-3 (Cell Signaling, #9661, 1:200), rabbit anti-Uteroglobin (Abcam, #ab40873, 1:500). Isotype control rabbit (Biolegend, #910801), rat (Biolegend, # 400602) and mouse (Biolegend, #401402) antibodies were used. Secondary antibodies were diluted in 2% donkey serum, 10% DMSO and 0.5% Triton-X-100 in PBS at 1:1000. All the following secondary antibodies were purchased from Invitrogen unless stated otherwise. Donkey anti-rabbit Alexa Fluor 568 (#A10042), donkey anti-mouse Alexa Fluor 568 (#A10037), donkey anti-rabbit Alexa Fluor 647 (#A31573), donkey anti-rat Alexa Fluor 647 (Jackson ImmunoResearch, #712-605-153), donkey anti-goat Alexa Fluor 647 (#A21447), donkey anti-rabbit Alexa Fluor 790 (#A11374), donkey anti-rat Alexa Fluor 790 (Jackson ImmunoResearch, #712-655-153;).

### Histopathology

Tissues from infected animals sacrificed at day 1-7 and day 14 as well as non-infected hamsters sacrificed at day 7 were included in the study (n=3 per day). The left lung lobe was carefully removed, immersion-fixed in 10% neutral-buffered formalin, paraffin-embedded, and 2–3-μm sections were stained with hematoxylin and eosin (HE) for light microscopy examination. Consecutive sections were processed for immunohistochemistry (IHC), conventional azan and Prussian blue staining (details given in Suppl. Table 3). Briefly for IHC, sections were mounted on adhesive glass slides, dewaxed in xylene, followed by rehydration in descending graded alcohols. Endogenous peroxidase was quenched with 3 % hydrogen peroxide in distilled water for 10 minutes at room temperature. Antigen retrieval was performed and nonspecific antibody binding was blocked by pure goat normal serum for 30 minutes at room temperature. Immunolabeling was visualized by 3-amino-9-ethylcarbazole substrate (AEC, Dako, Agilent, Santa Clara, CA, USA), producing a red-brown signal and sections were counter-stained with Mayer’s hematoxylin. As positive controls for staining specificity, archived mouse spleen tissue section (AIF1) and archived SARS-CoV2-infected hamster lung tissue section (TTF1) were included in each run. As a negative control, a normal rabbit serum (AIF1, TTF1, SARS-NP) or an irrelevant antibody was used. All slides were scanned using Hamamatsu S60 scanner (Hamamatsu Photonics, K.K. Japan). NDPview.2 plus software (Version 2.8.24, Hamamatsu Photonics) was used for evaluation.

Evaluation and interpretation of histopathologic findings were performed by a board-certified pathologist (DiplECVP) in a masked fashion using the post-examination masking method (Ref: Meyerholz DK, ILAR J. 2018;59:13-17). A detailed semiquantitative, severity-based, ordinal scoring was applied on HE stained sections (details given in Suppl. Table 2). The sum of individual criteria resulted in i) vascular lesion, ii) inflammation, iii) necrosis, and iv) regeneration scores for each individual animal. The distribution of SARS NP protein was recorded in main and distal bronchi as well alveolar epithelium on ordinal scores using the tiers 0 = no antigen, 1 = rare, <5% per slide; 2 = multifocal, 6%–40% affected; 3 = coalescing, 41%–80% affected; 4 = diffuse, >80% affected.

Quantitative image analysis was performed using HALO software (Version: 3.2.1851.439, Indica Labs) with Multiplex IHC v3.0.4 (TTF1) and Area Quantification v2.1.11 module (AIF1) on the left main lung lobe from each animal. Slides were annotated to exclude glass and red blood cells. Manual inspection of all images was performed on each sample to ensure that the annotations were accurate.

### Tissue Optical Clearing and Immunolabelling

The hamster right caudal lung lobes were fixed for at least 21 days in 4% PFA before transfer to the BSL2 laboratory and then washed 3x in PBS/0.02% NaN_3_ daily for 3 days. After that, the lungs were cut into 350 µM sections using the vibratome (VT1200S, Leica Biosystems, Germany), and stored in PBS/0.02% NaN_3_ at 4°C until used. The immunolabelling and clearing protocols were performed according to [60, 61] with minor modifications. For negative controls, isotype and mock antibody-free staining was used. All following steps were performed with shaking at 120 rpm using a temperature controlled orbital shaker (New Brunswick Innova 42R, Eppendorf, Germany). Briefly the lung slices were dehydrated in a methanol gradient diluted in distilled water (v/v = 50%,80%, and 100%) for 1 hour each, and in the 100% step the methanol was replaced after 30 minutes of incubation. Then, the samples were bleached overnight in 100% methanol containing 5% hydrogen peroxide. The following day, the samples were rehydrated in methanol (80% and 50%) for 1 hour each and for the 50% methanol step, the solution was replaced after 30 minutes of incubation. After that, the samples were washed for 20 minutes 3x in PBS at room temperature. Then the samples were further washed in 0.2% Triton X-100/PBS solution for 1 hour twice at 37 °C as a pre-permeabilization step. For permeabilization, the samples were transferred to 0.2% Triton X-100/20% DMSO/0.3 M glycine/PBS, for 48 hours at 37°C. After permeabilization, the samples were blocked in 10% Donkey serum/10% DMSO/0.5% Triton X-100/PBS, at 37°C for 48 hours. Then, the primary antibodies were applied for 3 days and washed in 2% BSA, and 0.5% Triton X-100 in 1x PBS for 3 hours with 4 buffer changes in the first hour (i.e. every 15 minutes) and then 4 buffer changes over the remaining 2 hours (i.e. every 30 minutes). Following the washing step, the secondary antibody was applied for 3 days and washed as previously mentioned in the primary antibody washing step.

The samples were embedded in 1% phytagel prepared in PBS in Cryomold embedding dishes (Laborversand, CMM Intermediate 4566). The samples were dehydrated in an ethanol gradient prepared in Aqua ad injectabilia (pH 9-9.5; Aponeo 08609338) (v/v = 30, 50, 70 and 100%) for >6 hour each and the 100% step twice. Following the ethanol dehydration step, the ethyl cinnamate (ECi) was added and exchanged after 6 hours and the samples were further incubated until clear for imaging.

### Light Sheet Fluorescence Microscopy

Volumetric imaging of lung sections was performed using Mylteny Biotech LaVision UltraMicroscope II with an Andor Zyla 5.5 sCMOS Camera, an Olympus MVX-10 Zoom Body (magnification range: 0.63–6.3x), and an Olympus MVPLAPO 2x objective (NA = 0.5). Z-stacks were recorded in 16-bit TIFF image format using LaVision Bio-Tec ImSpector Software (v7.0.127.0) with 2 μm step size either and 1x magnification for entire sample overview, or 4x for higher resolution imaging of selected regions. The light sheet was set to NA 0.156 with 100% light sheet width. Chromatic correction was applied for each fluorescence channel.

### 3D Image analysis

LSFM records were processed and quantified within Arivis Vision4D 4.0 (Arivis AG) software platform. The raw Tiff image stacks were converted to Sis format using Arivis SIS Converter 3.5.1 (Arivis AG). An analysis pipeline was implemented using Arivis Vision4D machine learning trainer module for image segmentation. To perform model training, eight object classes were manually annotated to identify the stromal tissue, blood vessels, viral infection and specific cellular markers. A 2D feature set which included intensity and edge parameters, and 20% probability threshold was applied to train the model. The training results were estimated using probability maps for each class. For image stack segmentation and classification, 3D object connectivity and object feature filters were applied. The resulting segmented image stacks were volumetrically rendered and object feature quantitative data were exported in table format.

### Confocal Microscopy

In order to analyze tissue sections by confocal laser scanning fluorescence microscopy, the samples were removed from phytagel and transferred into chambers created using a 3D printer, as previously described in [61]. Leica Stellaris 8 microscope equipped with HC PL APO 40x/1,10 W CORR CS2 objective was used to acquire 3D image stacks, which were processed and analyzed using Arivis Vision4D software.

### RNA sequence analysis

RNA was extracted from homogenized right caudal accessory lung lobes using Direct-zol RNA Miniprep Kit (Zymo research, Freiburg, Germany) following manufacturer’s instructions. The samples from three animals were pooled for further analysis.

All RNAseq analysis steps were performed using CLC Genomics workbench v22.0.2 (Qiagen, Hilden, Germany). Raw fastq reads were checked for quality and trimmed accordingly. Due to insufficient RNA depletion during library preparation, trimmed reads were mapped to 28S ribosomal RNA (LOC121137942) and unmapped reads only mapped to the golden hamster scaffold BCM_Maur_2.0. All data are accessible at the Gene Expression Omnibus (GEO) database, https://www.ncbi.nlm.nih.gov/geo (accession no. GSE225382). To analyse deregulation of specific pathways, mouse gene ontology annotation was employed to examine transcription levels of genes implicated in relevant biological processes.

Statistical analysis for gene expression was conducted on the mapped reads. Visualization was done with in-house R scripts using the tidy verse library and ComplexHeatmap

### Illumina RNA sequencing

The total RNA samples (1 µg each) were further purified with DNA-free RNA kit (Zymo Research). All of the samples were subjected to rRNA removal using the QIAseq FastSelect -rRNA HMR Kit (Qiagen) and cDNA synthesis using Superscript IV (ThermoFisher Scientific) in the presence of Actinomycin D following the recommendations of the manufacturer. The resulting libraries were sequenced with an Illumina NextSeq 550 instrument with a single end 84-nucleotide setting. A detailed protocol of RNA-Seq library preparation from hamster lungs was described previously [62].

### Statistical analysis

Statistical analysis was done using GraphPad Prism 9.3 software, with exception to RNAseq data. Three biological replicates were used for each experiment and per time point. Individual data points, as well as average with standard deviation is shown unless otherwise stated. For two-group analysis, unpaired two-tailed t tests was used. Two-way ANOVA method was used for multiple group analysis. Asterisks show significance levels: * p<= 0.05, ** p<=0.01.

## Acknowledgements

We thank Kathrin Müller, Sophienne Touihri, Robin Brandt, Gabriele Czerwinski, Silvia Schuparis for technical assistance (FLI).

Funding was received through the European Joint Programme One Health EJP COVRIN project funded under the European Union’s Horizon 2020 Research and Innovation Programme https://onehealthejp.eu/jip-covrin/ [Grant number 773830].

OB thanks German Academic Exchange Service (DAAD) for scholarship funding under German Egyptian Research Long-term Scholarship (GERLS) program.

We acknowledge Prof. Nikolai Simmons, Prof. Sven Hammerschmidt (University of Greifswald) and the DFG grant number 447143887 for providing Leica Stellaris 8 confocal microscope.

---

**Supplementary Figure 1.**
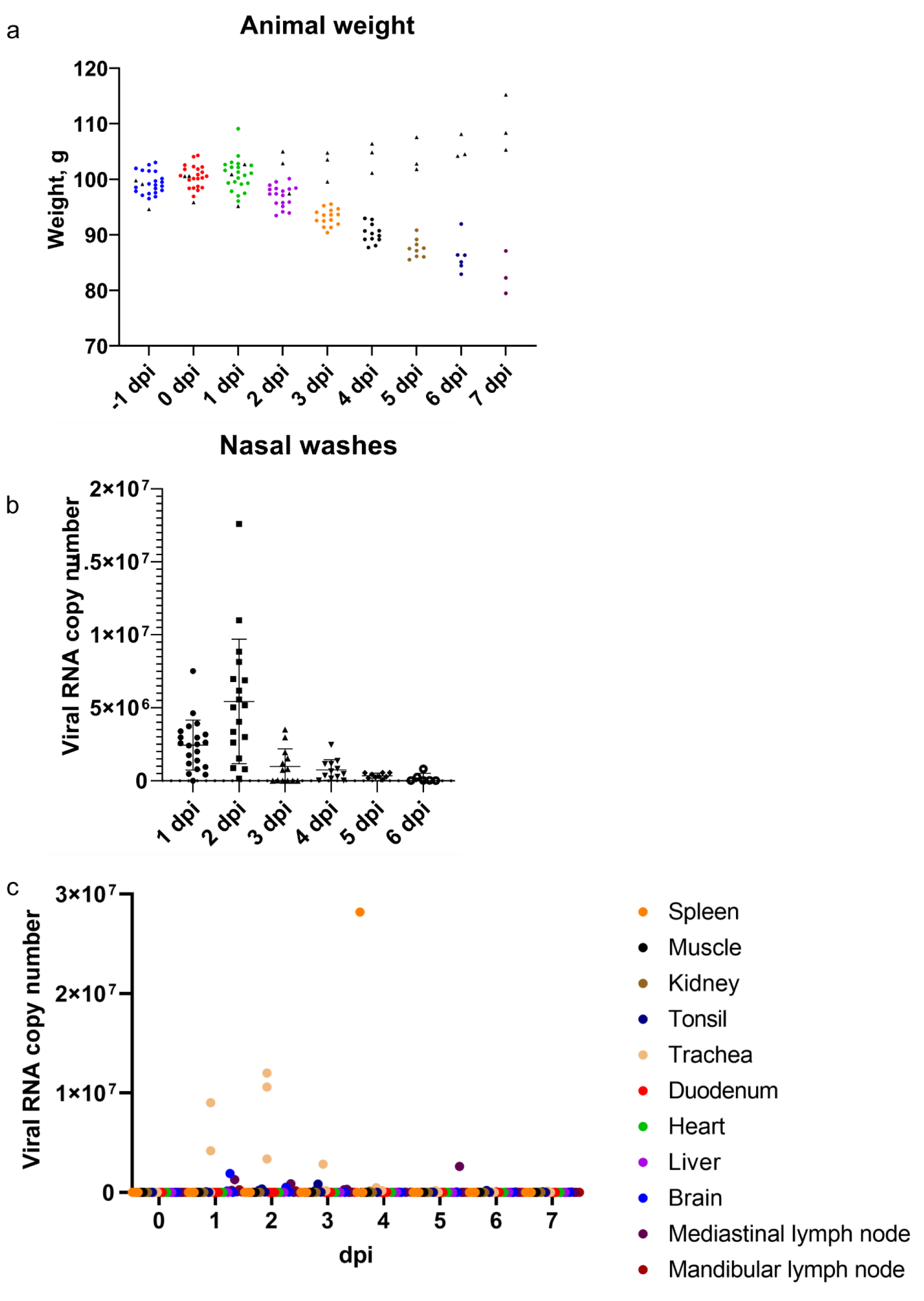
Animal weight and PCR analysis of viral RNA in nasal washes and different tissues. a) Animal weight measurements during the infection time-course. All animals were weighed one day prior infection (-1 dpi), on the day of infection (0 dpi) and on the following days. Control animal data is shown in black triangles. b) PCR analysis of viral RNA copy number (Log_10_) in nasal washes. c) PCR analysis of viral RNA copy number (Log10) in examined tissues. Each point in b and c represents data from a nasal wash or approximately 1.5-2 mm^3^ tissue sample, or the entire tissue in case of lymph nodes, of one animal.

**Supplementary Figure 2.**
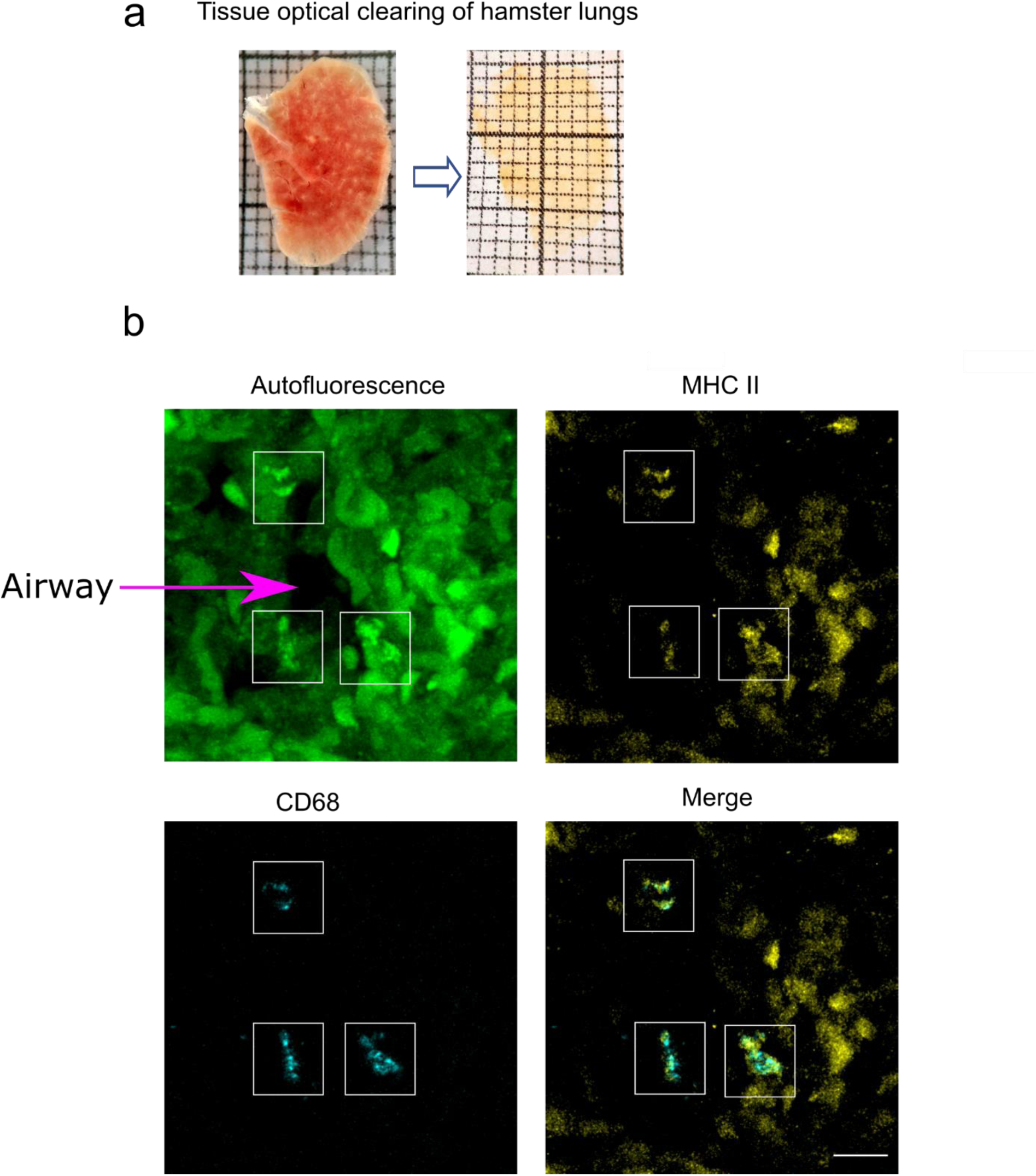
Tissue optical clearing and alveolar macrophage characterization in control lung tissues. a) Representative images of hamster lung tissue before (left) and after clearing (right) are shown b) Confocal imaging of control lungs showing alveolar macrophages. Maximum intensity projections of autofluorescence at 488 nm (green), CD68 (cyan), MHC II (yellow), and a merge of MHC II and CD68. White rectangles indicate location of CD68^+^, MHC II^+^ alveolar macrophages. Scale bar 10 μm.

**Supplementary Figure 3.**
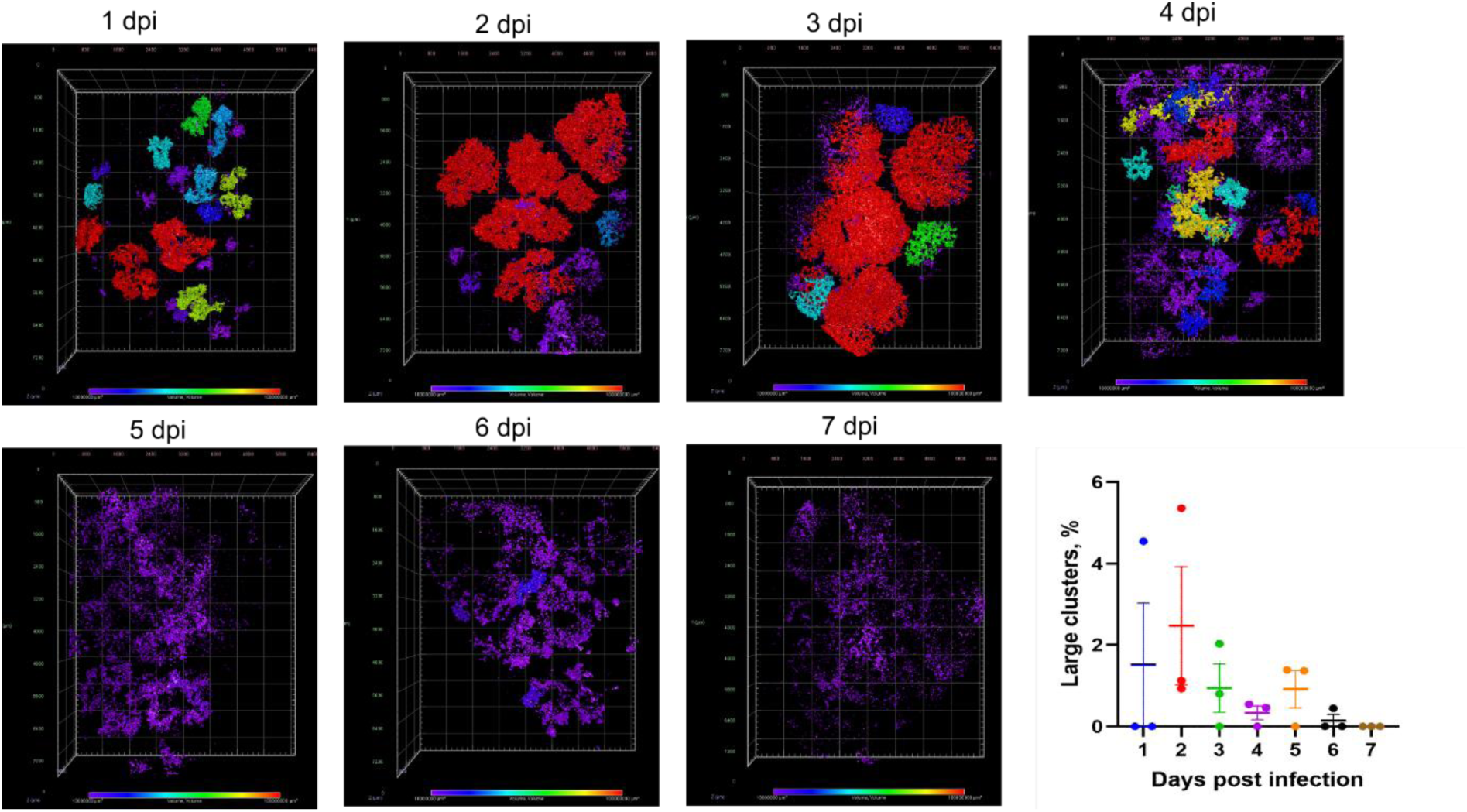
SARS-COV-2 node size distribution during the infection time-course. a) The NP fluorescence of samples from 1 to 7 dpi shown in Fig. 2a was classified according to the NP cluster size and displayed as 3D colour-coded heatmap. b) Distribution of large NP clusters shown as a fraction of total cluster number from 1 to 7 dpi.

**Supplementary Figure 4.**
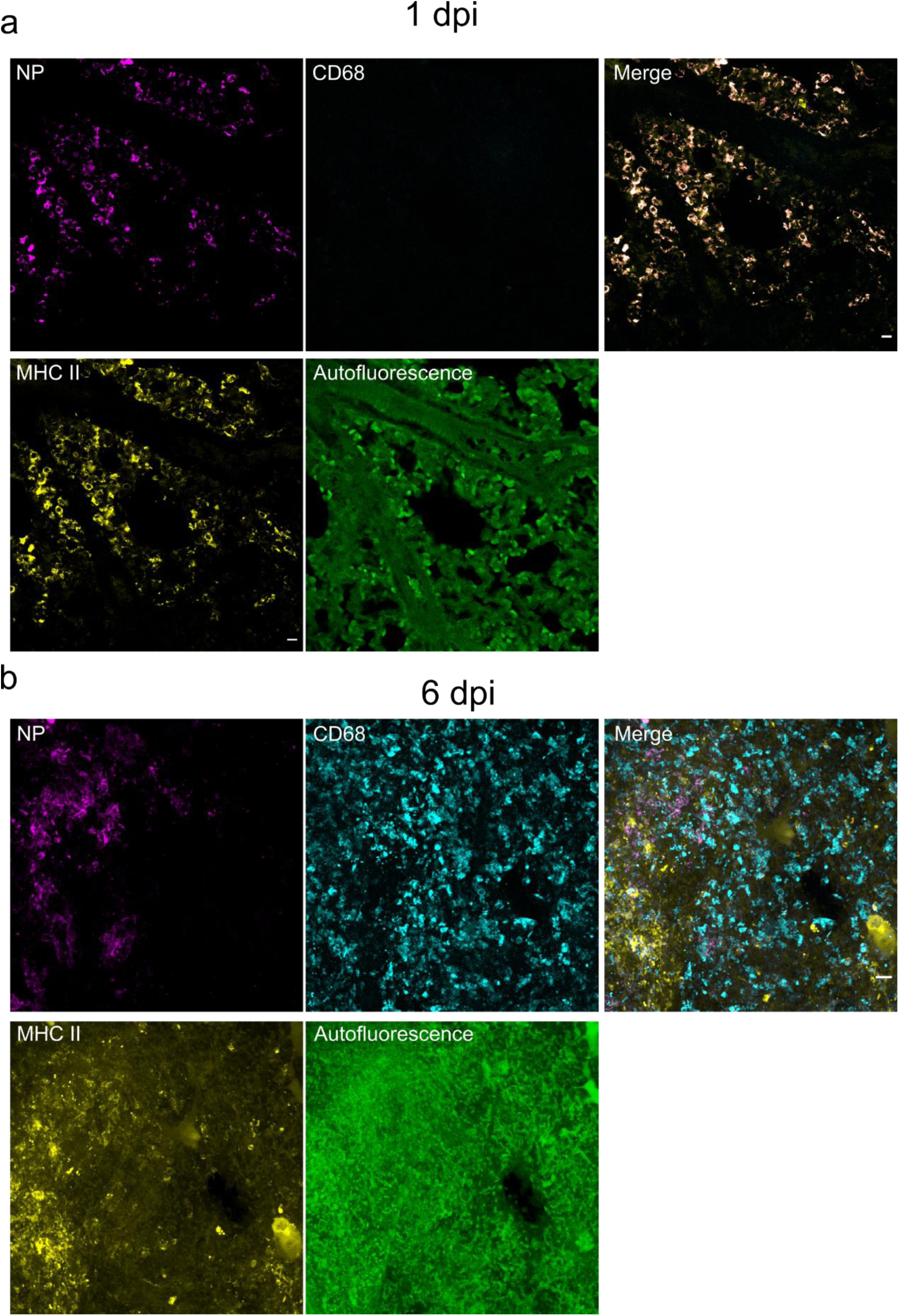
Confocal imaging of lung infection, MHC II and CD68 at 1 dpi (a) and 6 dpi (b). Maximum intensity projections of NP (magenta), CD68 (cyan), MHC II (yellow), autofluorescence at 488 nm (green). Scale bar 50 μm.

**Supplementary Figure 5.**
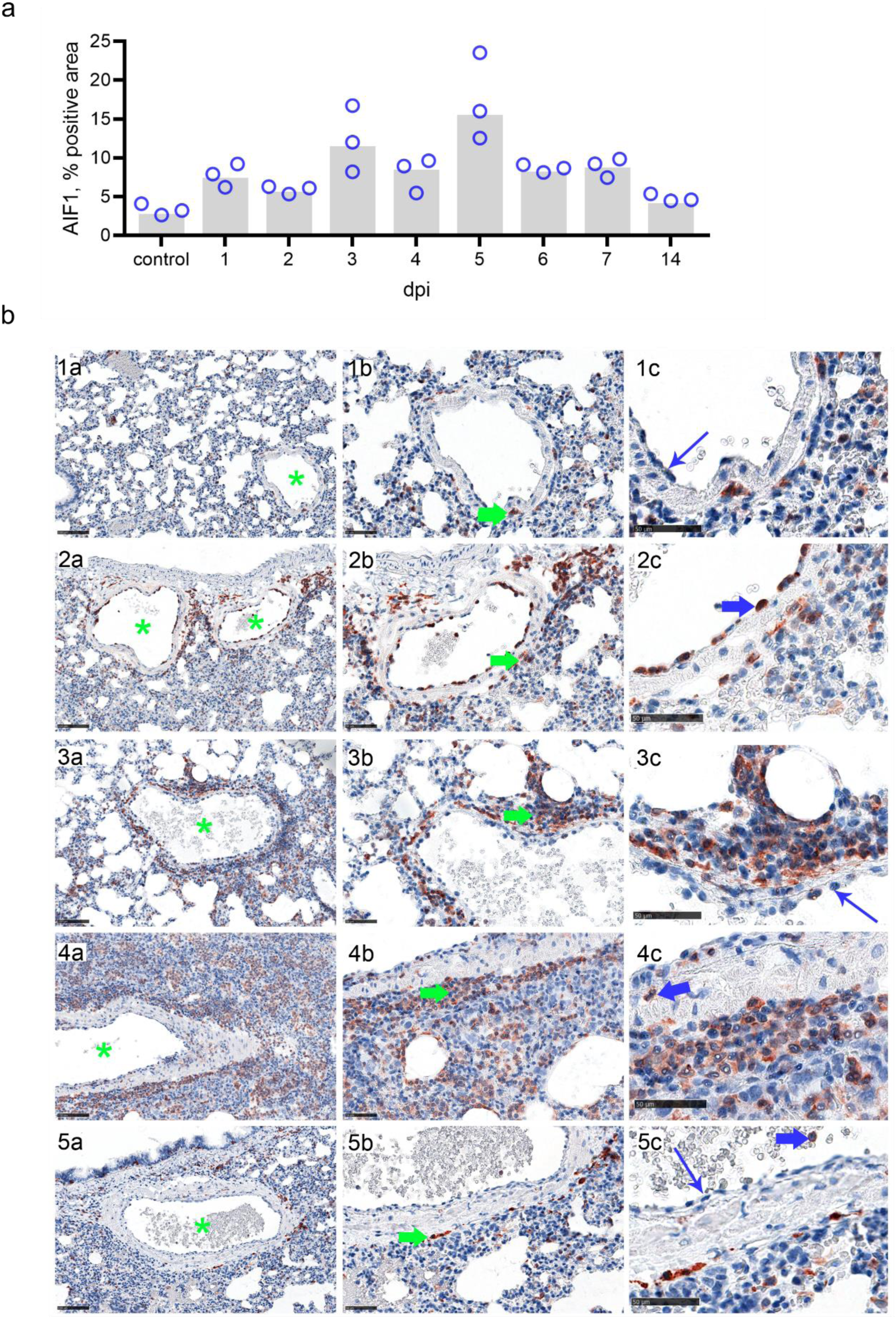
Spatiotemporal distribution of macrophages. (a) Immunohistochemistry (IHC) and quantitative 2D image analysis on AIF1 labelled lung slides for temporal evaluation. Waves of macrophage **influx peaking** on day 5 and almost reaching the baseline on day 14. HALO software (Version: 3.2.1851.439, Indica Labs), Area Quantification v2.1.11 module. Dots, relative AIF-positive area results for individual animals; bar, group median. **(b) Representative pictures showing IHC for AIF1 for spatial evaluation.** In control animals (**1**), Day 1 (**2**), day 5 (**3** less affected, **4** strongly affected area) and day 14 (**5**) post infection. Pictures showing an overview with perivascular (**a**, green asterisk indicating blood vessel lumen) and intra-alveolar macrophages (**a**, blue asterisk) with cytoplasmic AIF1 labelling. Image detail indicating perivascular macrophages (**b**, green arrow), image detail showing unlabelled endothelial cells (**1c, 3c, 5c**, blue slim arrow), and AIF1-positive monocytes/macrophages either intravascularly (**2c**, **5c** blue thick arrow) or transmigrating (**4c** blue thick arrow). Note that one day post infection, monocytes/macrophage constitute the main immune cell population involved in immune cell “rolling” (2c). Immunohistochemistry, AIF1 antigen, ABC Method, AEC chromogen (red), hematoxylin counterstain (blue), bar 100 µm (a) and 50 µm (b-c).

**Supplementary Figure 6.**
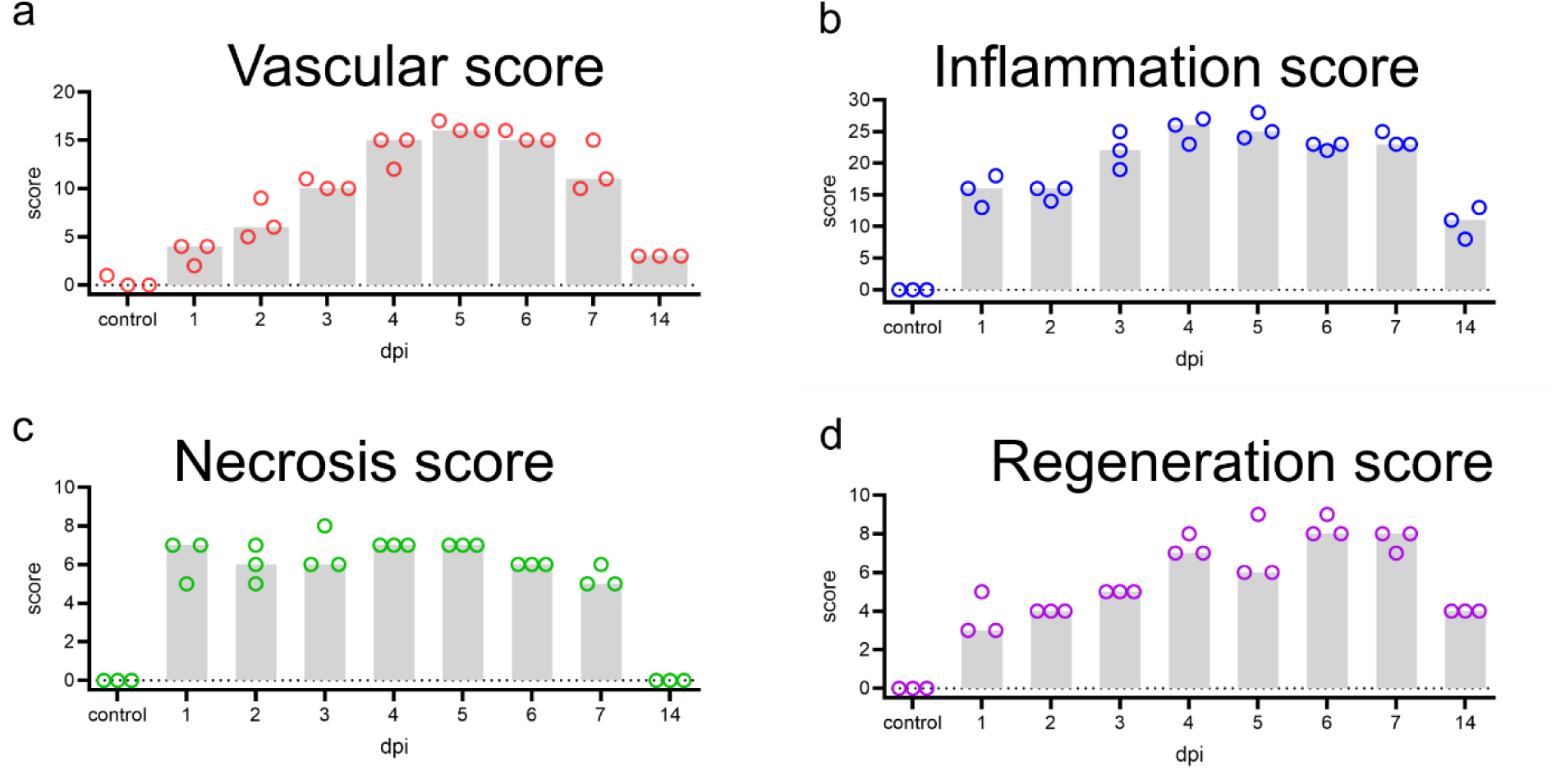
Detailed lung lesion scores. Inflammatory (a), vascular (b), necrosis (c) and regeneration (d) scoring was performed using scoring criteria described in Suppl. Table 2.

**Supplementary Figure 7.**
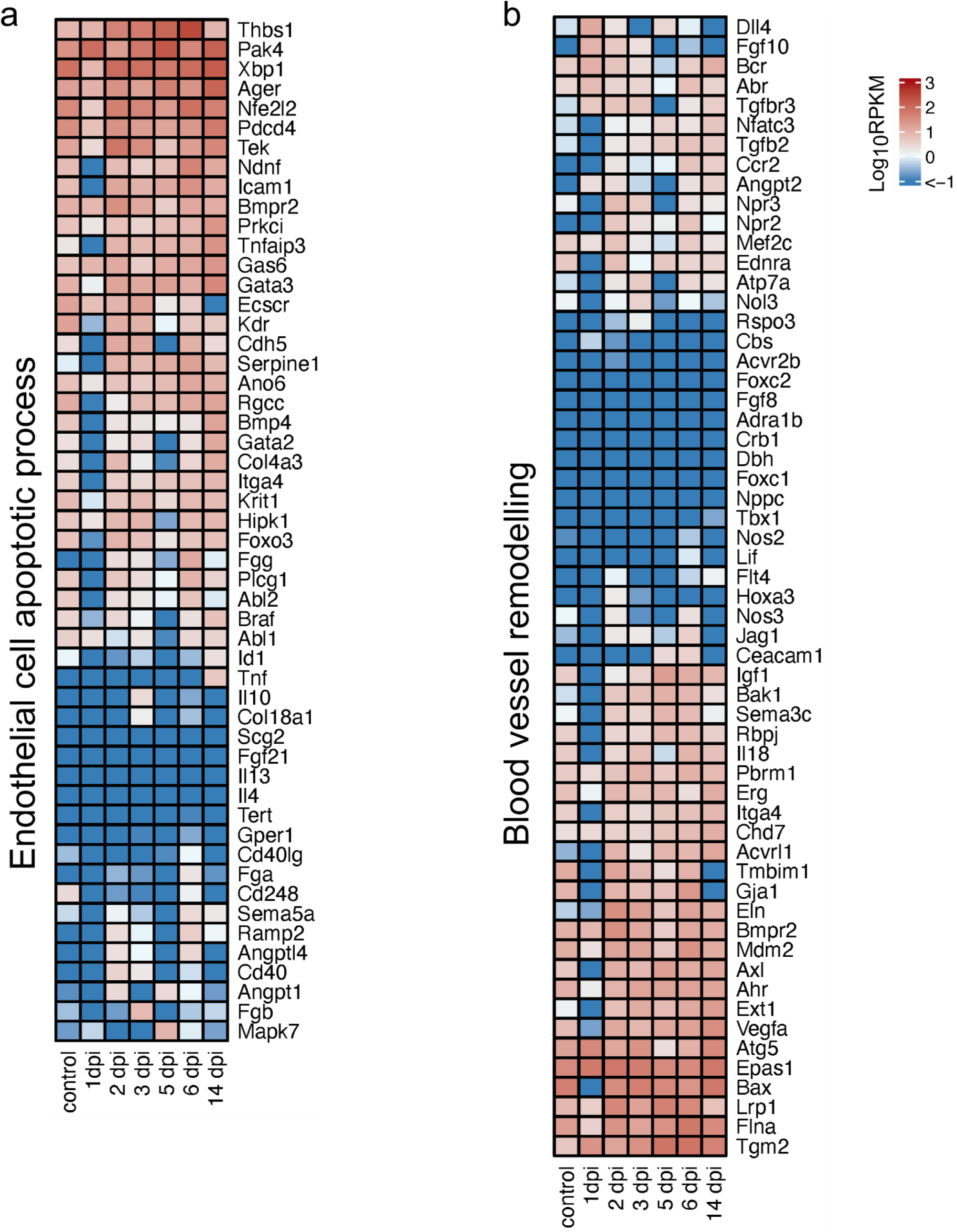
**Heat maps depicting scaled mRNA expression of endothelial apoptosis and vessel remodelling.**

**Supplementary Figure 8.**
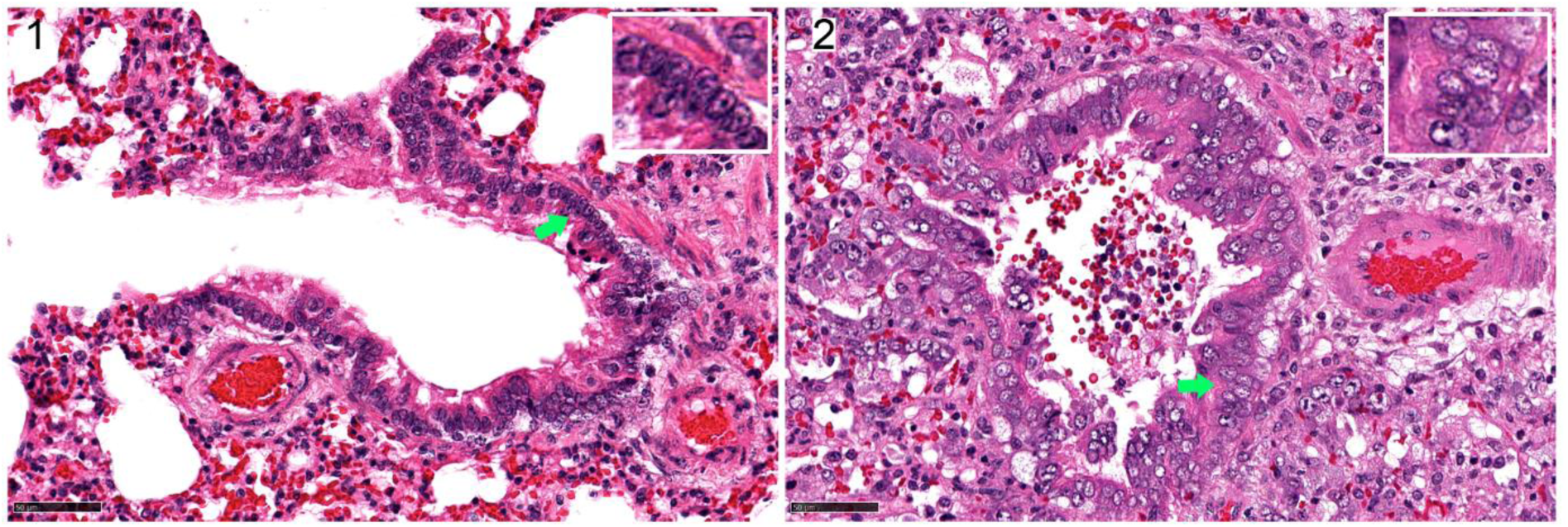
Regeneration of conducting airways. Control animals present with pseudostratified columnar epithelium, containing round to oval heterochromatic (inlay) nuclei in terminal airways (**1**). SARS-CoV-2 infection associated damage of the bronchial epithelium leads to regenerative hyperplasia and hypertrophy starting on day 2, here exemplarily shown on day 6 (**2**). Cells exhibit euchromatic (inlay) nuclei. Hematoxylin eosin stain, bar 50 µm.

**Supplementary Movie 1. 3D visualization of raw and processed LSFM records.** A lung section at 5 dpi showing unprocessed and processed views with different classes of cells and objects (NP, MHC II, MDM, tissue and blood vessels) identified by machine learning based segmentation. Note MDM clustering around major blood vessels.

**Supplementary Table 1.**
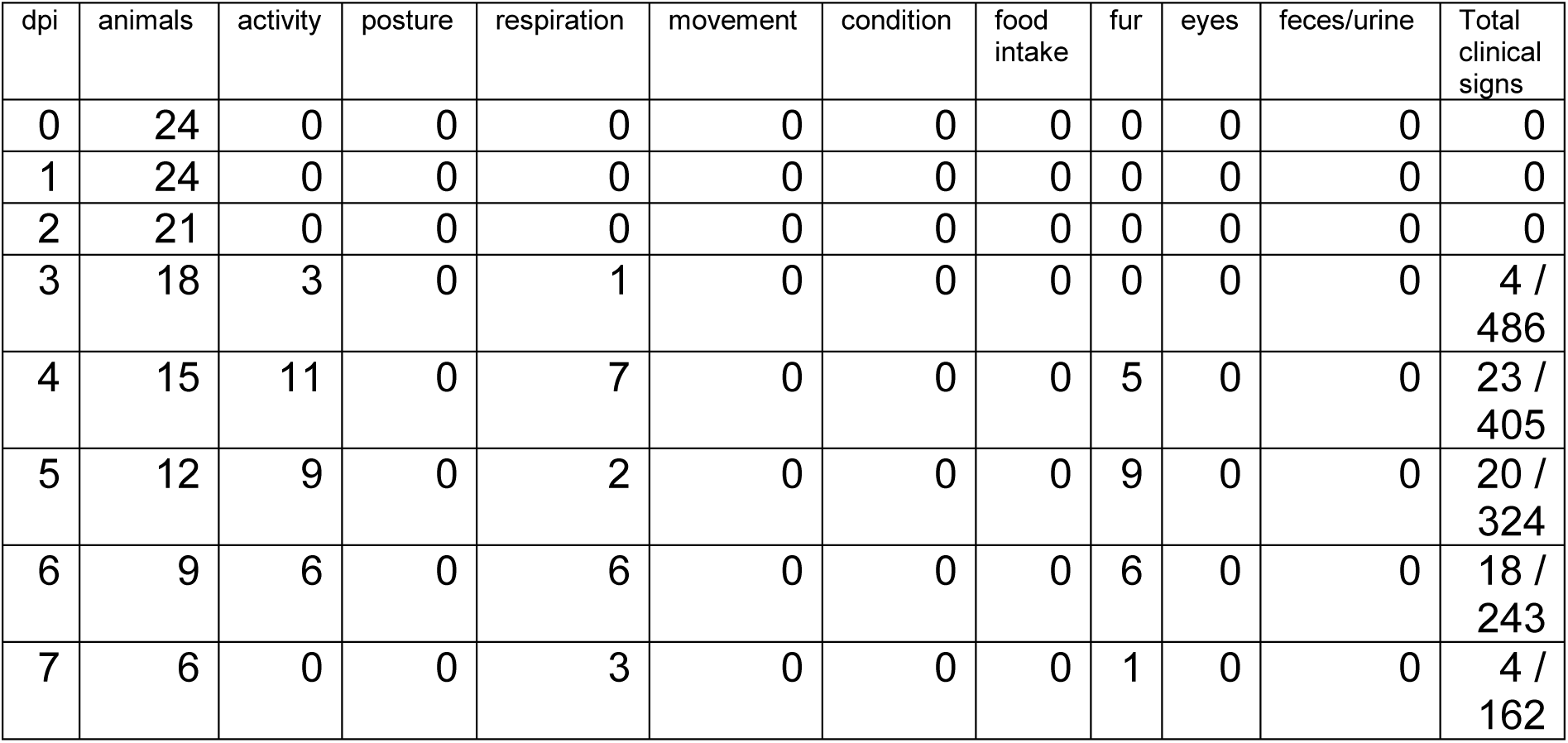
Animal observation scoring. For each criterion, a maximum of 3 points was possible per animal (i.e. 18 animals: maximum score of 54).

**Supplementary Table 2.**
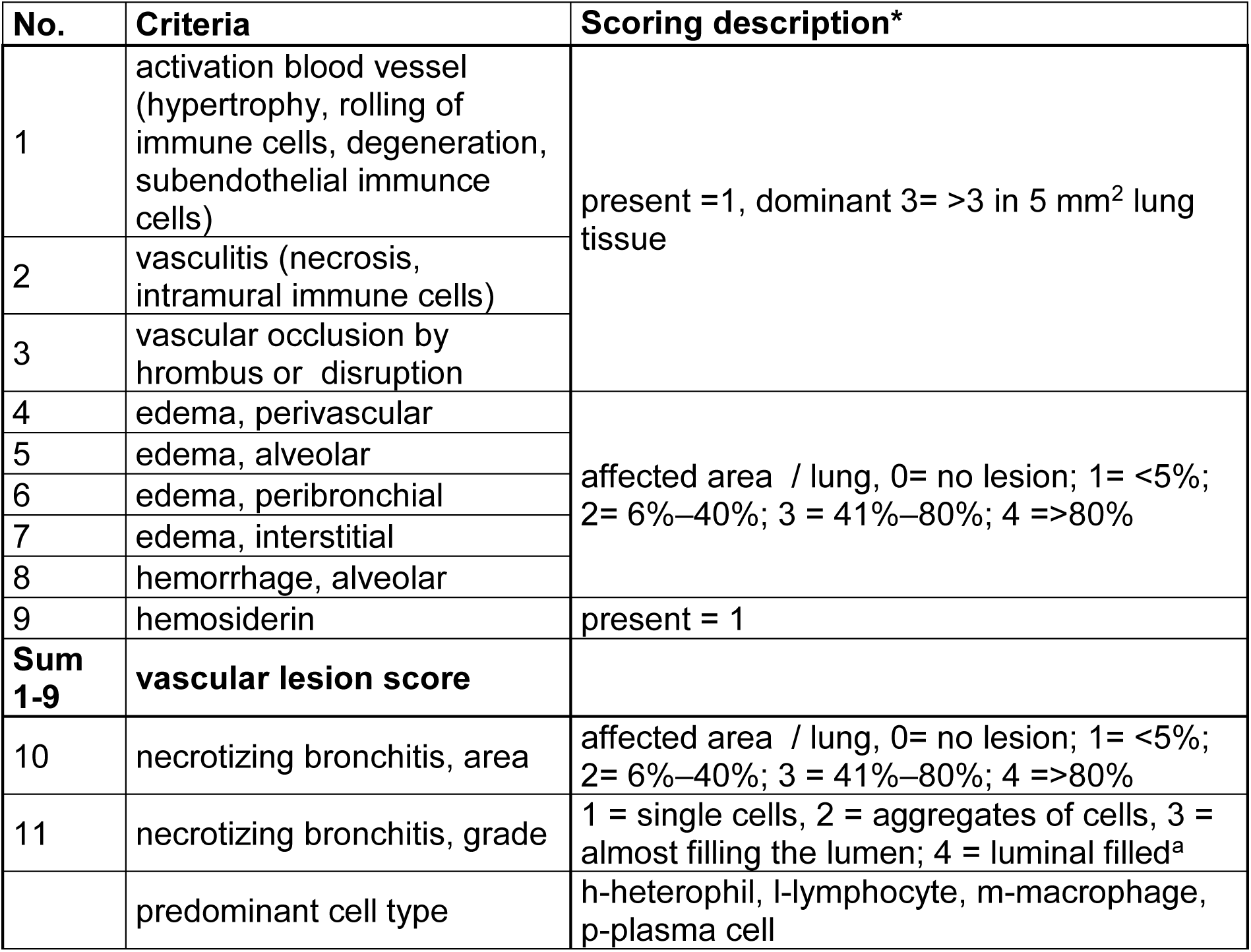
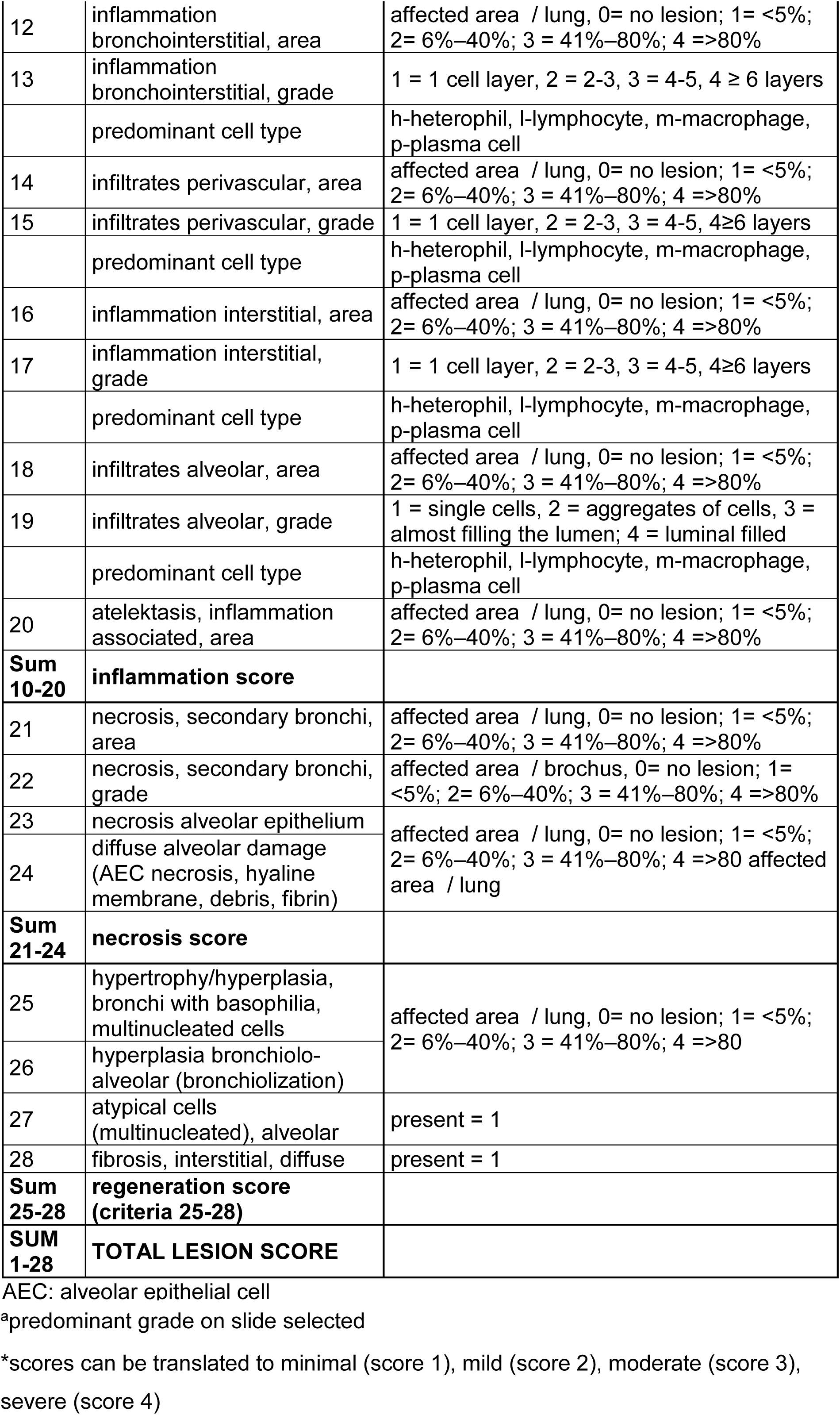
Histological scoring criteria

**Supplementary Table 3.** Primary antibodies used for IHC, applications, special staining and image analysis including measurement parameters

Immunohistochemistry
| Marker | Antibody | Pre-treatment | Secondary reagents | Measurement parameter |
| --- | --- | --- | --- | --- |
| SARS-CoV NP | Rabbit anti-SARS-CoV NP (Novus Biologicals, #NB100-56576), 1:200, ON | HIER, Citrate buffer pH 6.0, for 20 min | Anti-rabbit IgG Biotinylated (Vector Laboratories), 1:200, 30 min. RT; and ABC Kit Vectastain Elite PK 6100, 30 min, RT (Vector Laboratories) | Positive area per total tissue area |
| AIF1 | Rabbit anti-Iba1 (FUJIFILM Wako Pure Chemical Corporation, #019-19741), 1:800, ON | HIER, Citrate buffer pH 6.0, for 20 min | Dako EnVision+ System- HRP Labelled Polymer Anti-rabbit, 30 min, RT | Positive area per total tissue area |
| TTF-1 | Rabbit anti-TTF1 (abcam, ab#76013), 1:100, ON | HIER, Citrate buffer pH 6.0, for 20 min | Anti-rabbit IgG Biotinylated, 1:200, 30 min. RT; and ABC Kit, 30 min, RT | Positive cells per total tissue area |

Special staining
| Name | Kit / Chemicals | Measurement parameter |
| --- | --- | --- |
| Azan stain | Heidenhain's AZAN trichrome stain kit, following user manual (MORPHISTO, #12079 00250) | Positive area per total tissue area |
| Prussian blue | Hydrochloric Acid Solution, Potassium ferrocyanide solution, Nuclear fast red-aluminum sulfate solution | 1 = 1-3 foci<br>2 = >3 foci<br>3 = confluent<br>4 diffuse |

Abbreviations: IHC, immunohistochemistry; PAX, paired box protein; CD, cluster of differentiation; AIF1, ionized calcium-binding adapter molecule 1; HIER, Heat induced
epitope retrieval; EDTA, Ethylenediamine tetraacetic acid; RT, room temperature; ON, overnight at 4°C

